# Patient-derived tau-seeded human neuronal chimeras recapitulate mature Alzheimer’s tau pathology and uncover human-specific neuronal vulnerability

**DOI:** 10.64898/2026.07.06.736894

**Authors:** Yanru Ji, Purba Mandal, Jiahui Liu, Wen Hu, Bryce David Colón, Changlin Wan, Raymond W Pan, Dongkai Guo, Lijia Zhang, Xingjian Gu, Yang Yang, Kun Huang, Riyi Shi, Chao Qi, Chongli Yuan, Jean-Christophe Rochet, Priyanka Baloni, Fei Liu, Ranjie Xu

**Author notes:** Address correspondence to: Ranjie Xu, Ph.D. Assistant Professor, Department of Basic Medical Sciences Purdue University, 625 Harrison St., West Lafayette, IN 47907.

## Abstract

Tau pathology is a central hallmark of Alzheimer’s disease (AD) and strongly correlates with cognitive decline, yet the development of tau-targeted therapies has been hindered by the inability of existing models to capture human-specific disease features, particularly the mature forms of AD tau pathology. Moreover, species-specific mechanisms underlying tau pathology remain poorly understood. Here, we establish a patient-derived tau–seeded human neuronal chimera model by transplanting human pluripotent stem cell (hPSC)-derived neural progenitor cells into neonatal mouse brains, followed by intracerebral injection of pathological tau seeds from postmortem AD brains. Human neurons matured *in vivo* and recapitulated adult human tau features, including all six isoforms with an approximately 1:1 3R:4R ratio. Upon seeding, aged human neurons without FTD mutations faithfully developed robust, mature AD tau pathology, including neurofibrillary tangles (NFTs) and neuropil threads composed of paired helical filaments (PHFs) and straight filaments (SFs) containing 3R and 4R tau, closely mirroring advanced-stage AD. This pathology accumulated and spread across anatomically connected regions, accompanied by neurodegeneration, elevated plasma pTau-217, and memory deficits. Strikingly, tau pathology was largely restricted to human neurons, revealing a pronounced human-specific vulnerability. Mechanistically, snRNA-seq showed that human neurons exhibited higher basal expression of tau-uptake genes and widespread synaptic suppression following tau exposure, whereas mouse neurons remained transcriptomically resilient. Finally, the familial AD mutation *PSEN2* N141I exacerbated tau pathology and synaptic loss in human neurons. Together, this model recapitulates the molecular, structural, and functional hallmarks of mature AD tau pathology in human neurons *in vivo* and reveals intrinsic, species-specific vulnerability, providing a human-relevant *in vivo* platform for mechanistic studies and therapeutic development.

## Introduction

Tau pathology, characterized by the abnormal aggregation of hyperphosphorylated microtubule-associated protein tau, is a central hallmark of Alzheimer’s disease (AD) and strongly correlates with cognitive decline^1–3^. Despite promising preclinical results, most tau-targeted therapies have failed in clinical trials, highlighting the need for improved mechanistic understanding and more translational models^4, 5^. In AD brains, pathological tau forms paired helical filaments (PHFs) and straight filaments (SFs) that assemble into neurofibrillary tangles (NFTs) and neuropil threads, and propagate across interconnected brain regions in a stereotypical pattern^6–9^. These structures represent mature forms of tau pathology and are closely associated with neurodegeneration and disease severity^5, 6, 10^. However, the mechanisms underlying tau pathology and its contribution to disease progression remain poorly understood^5, 6^, significantly hindered by a lack of models that faithfully recapitulate these mature pathological features. Furthermore, whether human neurons exhibit intrinsic vulnerability and distinct mechanistic responses to tau pathology remains a critical, unresolved question with direct implications for therapeutic development.

Existing tau models have substantially advanced the field but are limited in recapitulating human-specific features of AD tau pathology. A major limitation is that most transgenic mice rely on frontotemporal dementia (FTD)-associated *MAPT* mutations (e.g., P301L/S) to induce tau pathology, yet such mutations are absent in AD^11–13^. In addition, tau isoform composition differs between species^14–16^. In adult humans, six tau isoforms are expressed with an approximately balanced ratio of 3-repeat (3R) and 4-repeat (4R) tau, whereas adult rodents predominantly express 4R tau^14–18^. Given that perturbations of the 3R:4R ratio contribute to tauopathy pathogenesis and both isoforms are involved in AD, faithfully recapitulating human tau biology in rodents is inherently challenging^14, 18, 19^.

Beyond these limitations, fundamental genomic and physiological divergences between human and animal neurons present a critical barrier to modeling human-specific disease mechanisms underlying tau pathology^12,20,21^. These include species-specific differences in gene regulation, alternative splicing, and neuronal excitability, which may have profound impacts on disease processes^12,20,21^. Importantly, the lack of systems that allow direct comparison between human and animal neurons *in vivo* has limited the identification of intrinsic, species-specific vulnerabilities to tau pathology. Defining these mechanisms is essential for understanding disease progression and developing effective therapies.

Although existing human models reproduce important aspects of AD tau pathology, challenges remain in simultaneously recapitulating mature tau pathology together with neuronal aging, anatomical tau spreading, plasma biomarker changes, and cognitive dysfunction in human neurons^22–27^. Specifically, these models rarely develop AD-like NFTs or PHFs characteristics of advanced-stage human AD^22–26^. Thus, there is an urgent need for translational models that faithfully reproduce mature AD tau pathology in human neurons and uncover human-specific disease mechanisms.

To bridge this gap, we developed a patient-derived tau–seeded human neuronal chimera model by transplanting hPSC-derived neural progenitor cells (NPCs) into neonatal mouse brains, followed by intracerebral injection of pathological tau seeds from postmortem AD brains in adulthood. In this system, human neurons are widely distributed, mature *in vivo*, and recapitulate adult human tau features. Upon seeding, human neurons develop robust, mature AD tau pathology that propagates across interconnected brain regions. Notably, this model enables direct comparison between human and mouse neurons under identical *in vivo* conditions and reveals a human-specific neuronal vulnerability to tau pathology. Mechanistically, we identify distinct transcriptional responses in human neurons associated with tau uptake and synaptic dysfunction. Together, this work establishes a human-relevant *in vivo* platform to interrogate species-specific disease mechanisms and provides new insights into intrinsic human neuronal vulnerability in AD.

## Results

### Development of a patient-derived tau-seeded human neuronal chimera model recapitulating adult human tau features and tau pathology *in vivo*

To develop a model that recapitulates mature AD tau pathology in human neurons without introducing FTD-associated mutations, we transplanted NPCs derived from various hPSC lines neonatally into Rag1-/- immunodeficient mice following our previous studies^28, 29^ (Supplementary **Table S1**, Supplementary **Fig. S1A**). Building on the established seeding activity of patient-derived pathological tau in rodents^30–32^, we hypothesized that the tau seeds would induce tau pathology in human neurons in chimeras. We stereotaxically injected soluble pathological tau (P-tau) extracted from postmortem AD brain into the hippocampus of chimeras at 4 months of age; control chimeras received PBS (**Fig. 1A**). Human cells were tracked using human-specific antibodies (hN, Ku80, or huNCAM; Supplementary **Fig. S1B**), and by 12 months, Ku80^+^ cells were broadly distributed across the hippocampus, cerebral cortex, striatum, midbrain, thalamus, and hypothalamus (Supplementary **Fig. S1C, D**). The engrafted NPCs differentiated neurons, astrocytes, and oligodendrocytes, with a small fraction being inhibitory neurons (Supplementary **Fig. S1E-J**).

**Figure 1.**
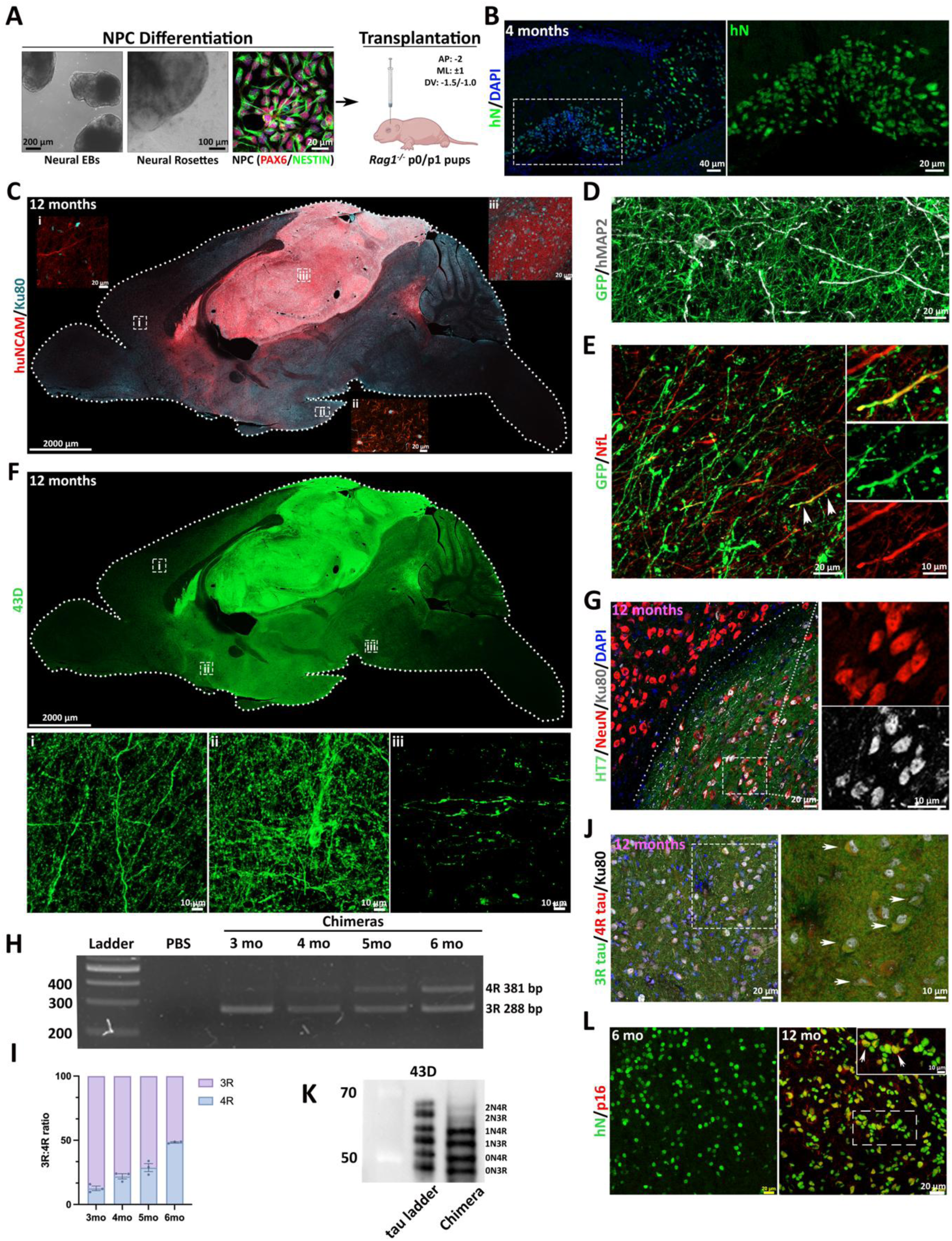
Development of a patient-derived tau-seeded human neuronal chimera model recapitulating adult human tau features and tau pathology *in vivo*. **(A)** Schematic of experimental design. Image was generated using BioRender. **(B)** Whole-brain image of 43D immunostaining showing widespread distribution of human tau within the host brain. 43D specifically labels human total tau. Insets indicate human tau protein distribution in **(Bi)** cortex, **(Bii)** striatum, and **(Biii)** midbrain. **(C)** Representative images of triple staining for HT7, NeuN, and Ku80, indicating that a large proportion of transplanted cells differentiate into neurons expressing tau. **(D)** Representative images of co-staining of 3R tau, 4R tau, and Ku80 showing that human neurons express both 3R and 4R tau. Arrows indicate human neurons exhibiting 3R- and 4R-tau colocalization. **(E)** Semi-quantitative PCR analysis of human 3R- and 4R-tau expression showing an age-dependent increase in 4R tau. **(F)** Quantification of the 3R:4R tau expression ratio (n = 3 per group). **(G)** Western blot of brain lysates from chimeras probed with 43D showing chimeras express six tau isoforms. **(H)** Representative AT8 immunostaining in hippocampus and adjacent cortex. No AT8 signal is detected in control chimeras (14-month-old). AT8⁺ cells in tau-seeded chimeras are indicated. Arrows indicate pre-tangle-like signals. Arrowheads indicate NFT-like signals. **(I)** Quantification of the number of AT8+ cells, and the area of the AT8+ neuropil threads across the hippocampus and cortex. tau_3 mpi, n = 3; tau_6 mpi, n = 4; tau_10 mpi, n = 6. Data are mean ± SEM; one-way ANOVA with Tuckey’s multiple comparisons test for the number of AT8^+^ cells, Welch’s ANOVA with Dunnett’s T3 multiple comparisons for AT8⁺ neuropil threads; *p < 0.05, **p < 0.01, ****p < 0.0001. N.D., not detected.

Human-specific tau features were examined in the chimeras. Human tau protein, detected by human-specific tau antibody 43D, exhibited near whole-brain distribution (**Fig. 1B**). Triple staining of HT7 (pan human tau), NeuN, and Ku80 confirmed that transplanted human cells differentiated into neurons expressing tau (**Fig. 1C**), containing both 3R- and 4R-tau isoforms (**Fig. 1D**). The maturation of tau isoform was examined by semi-quantitative PCR with human-specific primers^24^ (Supplementary **Table S2**). Minimal 4R tau was detected at 3 months; expression progressively increased to an approximately 1:1 3R:4R ratio by 6 months (**Fig. 1E, F**), indicating human progressive maturation of human neurons and tau in chimeric brains to adult-like features. Western blot analysis of chimeric brain homogenates using 43D revealed all six human tau isoforms (**Fig. 1G**). The chimeras showed long life span (> 24 months) and exhibited p16^+^ signals at 12 months (Supplementary **Fig. S1K**), suggesting that transplanted human cells not only mature but also exhibit aging-associated features *in vivo*.

The concentration and seeding activity of postmortem AD brain-derived P-tau were confirmed before inoculation (Supplementary **Fig. S2A, B, Materials and Methods**). To assess tau pathology, we collected chimeric brains at three different time points post P-tau injection (**Fig. 1A**) and examined tau hyperphosphorylation using AT8 (pSer202/pThr205) immunostaining. Control chimeras showed no AT8 signal at any time point (**Fig. 1H**; Supplementary **Fig. S2C**). In P-tau–injected chimeras, AT8-positive pretangle-like signals (puncta-like AT8 signal) and neuropil threads appeared as early as 4 months post-injection (mpi). By 6 mpi, neurofibrillary tangle (NFT)-like signals (high-intensity AT8 signal) and neuropil threads were also observed in the cerebral cortex (**Fig. 1H**). By 10 mpi, dense AT8-positive neuronal somata and processes occupied both hippocampus and cortex, forming numerous intraneuronal, NFT-like signals and abundant neuropil threads (**Fig. 1H**). AT8 co-localized with neuron marker (NeuN) only; no colocalization was identified with glial markers (GFAP, MBP) (Supplementary **Fig. S2D-F**). Quantification confirmed a significant, time-dependent increase in AT8-positive neurons and neuropil threads in the hippocampus and cortex (control: not detected, AT8-positive neurons: tau_4mpi: 0.431 ± 0.172, tau_6mpi: 1.440 ± 0.222, tau_10mpi: 2.632 ± 0.190; AT8-positive neuropil threads: tau_4mpi: 0.299±0.093, tau_6mpi: 2.114±0.277, tau_10mpi: 6.050±1.073) (**Fig. 1I**).

Taken together, these results demonstrate robust human tau distribution throughout the host brain, with adult human tau features accurately recapitulated. AD-patient-derived tau seeds induce progressive, AD-like tau hyperphosphorylation in chimeras, establishing this model as a potential platform for studying human-specific features of tau pathology.

### The chimeric brains reveal intrinsic human-specific neuronal vulnerability to tau pathology

To determine whether pathology arose from human or mouse neurons in chimeric brains, we co-stained pathological tau markers with various human markers. Clear colocalization was observed between Ku80 and the conformation-dependent tau antibody MC1 (**Fig. 2A**), as well as 43D and pT217-tau (**Fig. 2B**). Since Ku80 signal can be sequestered or lost in heavily tau-burdened neurons^33^, we additionally used GFP-labeled human cells and the human-specific mitochondrial marker hCOX IV. Triple staining for AT8, hCOX IV, and GFP confirmed robust colocalization of pathology with human neurons (**Fig. 2C**); notably, mitochondria appeared less abundant in heavily tau-burdened human neurons, suggesting impaired mitochondrial homeostasis under tau pathology. Quantification confirmed that the majority of AT8^+^ cells arose from human neurons at both time points (tau_6mpi: %Human neuron: 73.563 ± 2.381, %Mouse neuron: 26.437 ± 2.381; tau_10mpi: %Human neuron: 75.280±1.430, %Mouse neuron: 24.720±1.430) (**Fig. 2D**). The disproportionate burden of tau pathology in human neurons suggested that human neurons are more vulnerable to tau pathology than neighboring mouse neurons. Consistent with this, AT8 immunoreactivity was absent in the hippocampus or cortex of wild-type *Rag1^-/-^* mice 10 months after the inoculation of the same amount of tau seeds (Supplementary **Fig. S2G**). The absence of the AT8 signal at 7 days post-tau inoculation (Supplementary **Fig. S2H**) indicates templated conversion of endogenous tau, ruling out residual input seeds.

**Figure 2.**
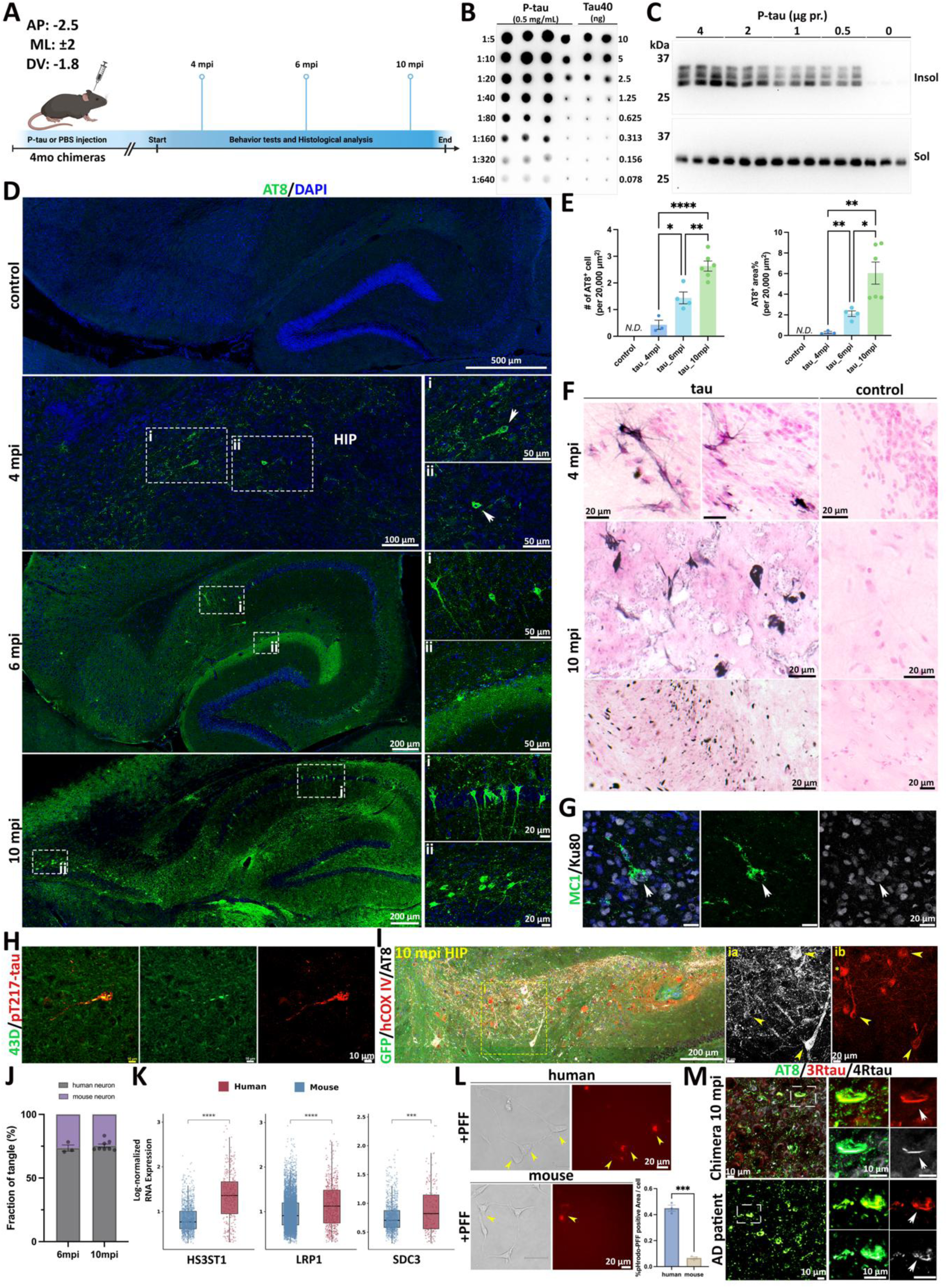
The chimeric brains reveal intrinsic human-specific neuronal vulnerability to tau pathology. **(A)** Representative image of MC1 co-stained with Ku80; arrow denotes a Ku80⁺ human neuron. **(B)** Representative 43D and pTau-217 double staining. Human tau signal partially overlaps with pTau-217, indicating 43D can serve as a marker for human tangle-bearing cells. **(C)** Representative triple immunostaining for AT8, hCOX IV, and GFP in hippocampus at 10 months post-tau injection. Insets show co-localization of AT8 and hCOX IV. Neurons with a heavier tau burden exhibit sparser hCOX IV puncta (arrowhead) relative to a comparatively pathology-free neuron (asterisk). **(D)** Quantification of the percentage of AT8^+^ cells in human versus mouse neurons at 6 mpi and 10 mpi. Images were taken from hippocampus and cerebral cortex. Various human markers were used to identify human neurons, including Ku80, 43D, GFP, and hCOX IV. Human neurons consistently contribute a greater proportion of total AT8^+^ cells. **(E)** Gene expression level of LRP1, HS3ST1, and SDC3 between human and mouse neurons. Data collected from cells showed gene expression level greater than 0, Wilcoxon t test; ***p < 0.001, ****p<0.0001. **(F, G)** Representative images showing K18 tau PFF uptake assay using embryonic stem cell-derived human neurons (left panel) and mouse neurons **(F).** Statistical results **(G)** indicate a significantly higher amount of K18 PFFs uptake by human neurons. Experiments were repeated 3 times. Data are mean ± SEM; unpaired two-tailed t test; **p < 0.01. Scale bars, as indicated in the panels.

To probe the basis of this species difference, we employed snRNA-seq to compare the expression of genes previously implicated in tau uptake and propagation (*LRP1*^34, 35^, *HS3ST1*^36, 37^, *SDC3*^38^, *HSPG*^39, 40^, and *SORL1*^41^) between human and mouse neurons (**Fig. 2E**). *LRP1*, *HS3ST1*, and *SDC3* were significantly higher in human neurons (human vs. mouse: *LRP1*: 1.144 ± 0.019 vs. 0.977 ± 0.005; *HS3ST1*: 1.330 ± 0.029 vs. 0.835 ± 0.009; *SDC3*: 0.901±0.029 vs. 0.758±0.008)(**Fig. 2E**), whereas *HSPG2* and *SORL1* showed no significant difference (human vs. mouse: *HSPG2*: 0.919±0.085 vs. 0.734±0.053; *SORL1*: 1.315±0.024 vs. 1.309±0.006) (Supplementary **Fig. S2I**). To functionally validate whether human neurons exhibit enhanced tau uptake ability, we performed a tau endocytosis assay using pHrodo-labeled K18 tau PFFs. Human neurons exhibited significantly greater endocytic uptake than mouse neurons under matched exposure conditions (human vs. mouse: 2.140±0.103 vs. 0.950±0.123) (**Fig. 2F, G**; Supplementary **Fig. S2J**), consistent with the gene expression data. Together, these results indicate that human neurons are intrinsically more vulnerable than mouse neurons to pathological tau seeds, likely in part due to a greater capacity for tau uptake.

### Patient-derived tau-seeded human neuronal chimeras recapitulate mature AD tau pathology with spread across interconnected brain regions

Beyond tau hyperphosphorylation, we next asked whether the chimeras would recapitulate mature AD tau pathology, characterized by NFTs and neuropil threads composed of PHFs and SFs, features of the advanced AD stage that are not reproduced in existing models. Gallyas silver staining revealed argyrophilic, insoluble filaments selectively in tau-seeded chimeras, indicating the formation of NFTs and neuropil threads (**Fig. 3A**, Supplementary **Fig. S3A**). To identify the tau isoform composition of NFTs in human neurons in tau chimeras, we triple-stained AT8 with 3R- and 4R-tau. Because mature adult mouse neurons express only 4R-tau, 3R reactivity within an inclusion marks a human origin. Our results indicate that both 3R- and 4R-tau contributed to AT8-postive inclusions in tau chimeras in human neurons, consistent with that observed in AD patient brains (**Fig. 3B**). Furthermore, we isolated sarkosyl-insoluble tau from seed-exposed chimeric brains following previous studies^42^ (Supplementary **Fig. S3B**). Negative-stain EM clearly revealed both SFs and PHFs closely resembling those observed in AD brain (**Fig. 3C**, Supplementary **Fig. S3C**), and immunogold EM with PHF-1 confirmed that these structures were composed of phosphorylated tau (**Fig. 3D**, Supplementary **Fig. S3D**). 43D and HT7 labeling demonstrated the human origin of insoluble aggregates (**Fig. 3E, F**, Supplementary **Fig. S3E, F**). To assess seeding capacity, sarkosyl-insoluble tau was applied to HEK293 tau biosensor cells as previously described^43^. Abundant intracellular fluorescent puncta were observed in both living cells and fixed cells, whereas vehicle-treated controls showed no detectable signal (**Fig. 3G, H**). Together, these results demonstrate that the model successfully recapitulates mature AD tau pathology.

**Figure 3.**
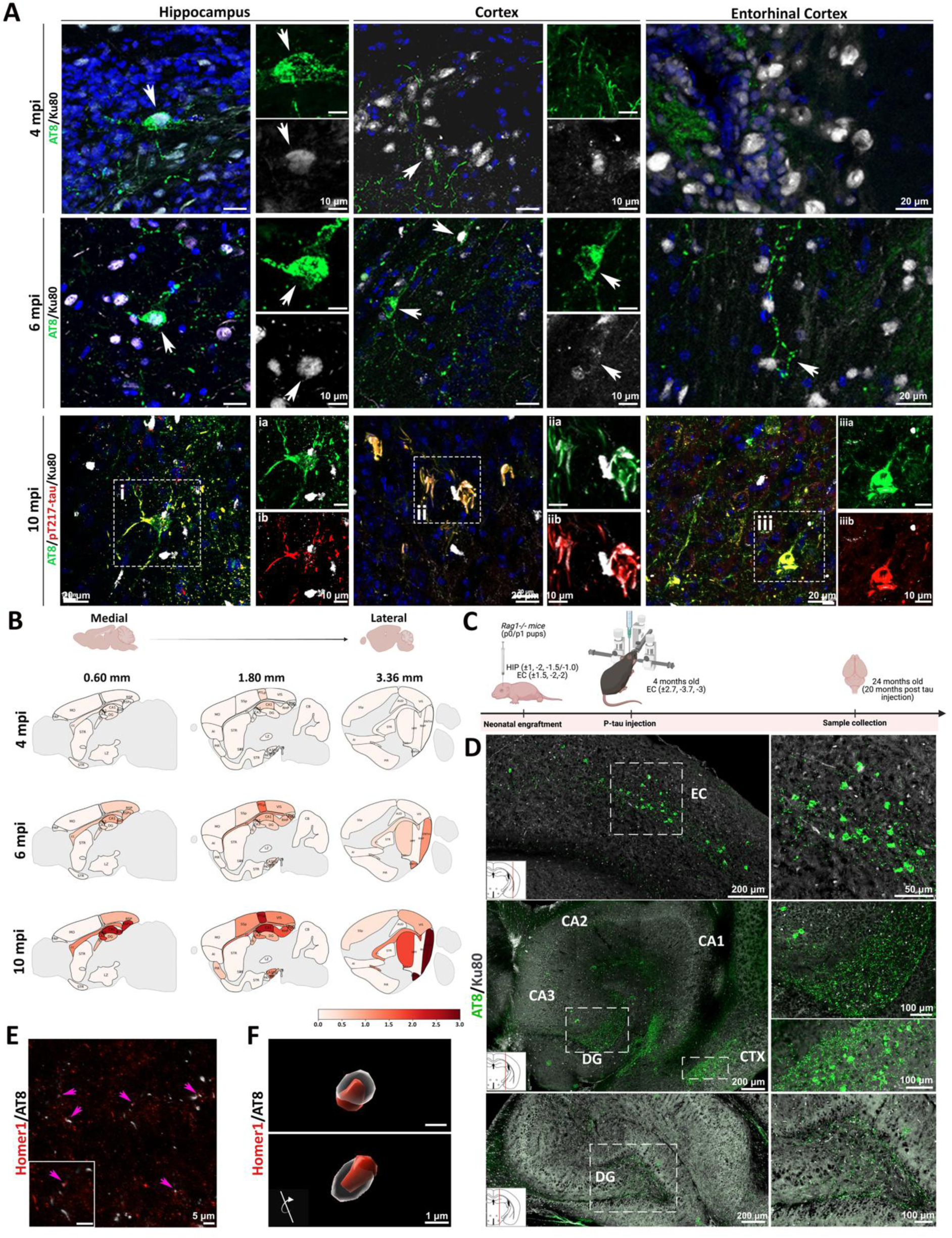
Patient-derived tau-seeded human neuronal chimeras recapitulate mature AD tau pathology with spread across interconnected brain regions. **(A)** Representative Gallyas silver staining in control, tau 4 mpi, and tau 10 mpi chimeras. NFTs and neuropil threads are observed only in tau-exposed chimeras. **(B)** Representative triple staining for AT8, 3R tau, and 4R tau in chimeras and AD patient samples. Arrows and insets demonstrate co-localization of both 3R and 4R tau with phospho-tau, indicating that chimeric tangles, like those in patients, contain both tau isoforms. Patient sample: mid frontal gyrus. Age: 73, Braak stage V. **(C)** Representative negative-stain EM images of isolated SI-tau. Both paired helical filaments (PHFs) and straight filaments (SFs) are detected from tau-seed–exposed chimeras. Asterisks indicate PHF structure. **(D–F)** Representative immunogold EM labeling of SI-tau from 14-month-old chimeras using **(D)** PHF1, **(E)** 43D, and **(F)** HT7. Arrows indicate gold particles attaching to fibrils. **(G, H)** HEK293T tau biosensor cells stably expressing the tau repeat domain with the P301S mutation fused to CFP or YFP were treated with vehicle or chimera-derived SI-tau. Representative images of living or fixed cells show no detectable YFP signal in vehicle controls, whereas YFP⁺ tau inclusions are observed following SI-tau exposure. **(H)** Quantification of YFP puncta–positive cells (n = 3). **(I)** Representative images of phospho-tau (AT8 or Thr217) immunostaining co-labeled with the human nuclear marker Ku80 across multiple brain regions at 4-, 6-, and 10-months post tau injection. Early, low-level pathology emerges at the injection site (hippocampus) by 4 mpi and progressively spreads to anatomically connected regions. Mature tangles are evident across three regions by 10 mpi (hippocampus, cortex, and entorhinal cortex). Arrows indicate tangle-bearing human neurons. **(J)** Heatmaps summarizing semi-quantitative AT8 pathology scores (0.0, no pathology; 3.0, severe pathology) at 4, 6, and 10 mpi. Three sagittal planes are shown (left to right). 4 mpi, n = 3; 6 mpi, n = 4; 10 mpi, n = 4. AI, agranular insular area; FRP, frontal pole; MO, somatomotor areas; SS, somatosensory areas; SSp, primary somatosensory area; RSP, retrosplenial area; RSPv, retrosplenial area, ventral part; RSPd, retrosplenial area, dorsal part; RHP, retrohippocampal region; HPF, hippocampal formation; CA1, field CA1; CA2, field CA2; CA3, field CA3; DG, dentate gyrus; fi, fiber tract; cc, corpus callosum; STR, striatum; PIR, piriform area; PA, posterior amygdala nucleus; CB, cerebellum; LZ, hypothalamatic lateral zone; PTLp, posterior parietal association areas; VIS, visual areas; AUD, auditory areas; ENTm, entorhinal area, medial part. **(K)** Representative AT8 and Homer1 double staining; arrows and the inset indicate overlapping AT8 and Homer1 signals. **(L)** Imaris-based 3D reconstruction of AT8 and Homer1 signals from different views. AT8⁺ puncta are closely opposed to Homer1⁺ postsynaptic structures. Scale bars, as indicated in the panels.

The seeding activity of tau is thought to drive the stereotyped spread of pathology from the entorhinal cortex (EC) to the hippocampus (HIP) and subsequently to the cerebral cortex (CTX) in AD patients^9, 44^. Thus, we examined AT8-positive pathology in Ku80^+^ human cells across EC, HIP, and CTX following hippocampal P-tau injection. At 4 mpi, pre-tangles were largely confined to the hippocampus, with faint labeling in the injection track within the posterior parietal association (PTLp). By 6 mpi, positive pathology had extended to the subiculum and visual cortex, with neuropil threads appearing in the EC. By 10 mpi, clear NFTs and neuropil threads were evident across all three regions (**Fig. 3I, J**), with the heaviest burden in CA1, subiculum, retrosplenial area, and EC, as summarized in a regional pathology heatmap (**Fig. 3J**). To more closely recapitulate the cortical origin of tau pathology in AD, a second cohort received NPC transplants into the HIP, CTX, and EC, followed by EC injection of P-tau at 4 months of age (Supplementary **Fig. S3G**). By 20 mpi, tau pathology had spread along connected circuits, including EC, dentate gyrus (DG), and CTX (Supplementary **Fig. S3G**). Co-staining of AT8 with the postsynaptic marker Homer1 revealed multiple colocalized puncta (**Fig. 3K, L**), suggesting synaptic involvement in transneuronal propagation. Collectively, these data demonstrate that patient-derived tau-seeded human neuronal chimeras faithfully recapitulate mature AD tau pathology, including the formation of NFTs and neuropil threads composed of PHFs and SFs containing 3R and 4R tau, as well as seeding competence, and propagation of pathology across interconnected brain regions.

### Tau pathology leads to neurodegeneration and glial responses in chimeric brains

Tau pathology correlates closely with neurodegeneration and cognitive decline^45–47^. To assess tau-associated neurodegeneration in our model, we examined human dendritic morphology and density using the human-specific dendritic marker hMAP2. In pathology-enriched regions, hMAP2 signal was markedly reduced, and many remaining dendrites appeared fragmented rather than forming continuous arbors (**Fig. 4A**). Quantification showed a time-dependent decline in hMAP2-positive area, with a trend toward reduction at 6 mpi and a further statistically significant decrease by 10 mpi, compared with controls (Control: 27.54±1.941; tau_6mpi: 14.67±3.172; tau_10mpi: 6.178±0.508) (**Fig. 4B**). Double labeling for hMAP2 and the postsynaptic marker Homer1 revealed a progressive loss of synaptic puncta on human dendrites that correlated with an increasing pathology burden (Control: 43.14 ± 2.481; tau_6mpi: 16.82± 0.978; tau_10mpi: 7.146 ± 1.685) (**Fig. 4C, D**).

**Figure 4.**
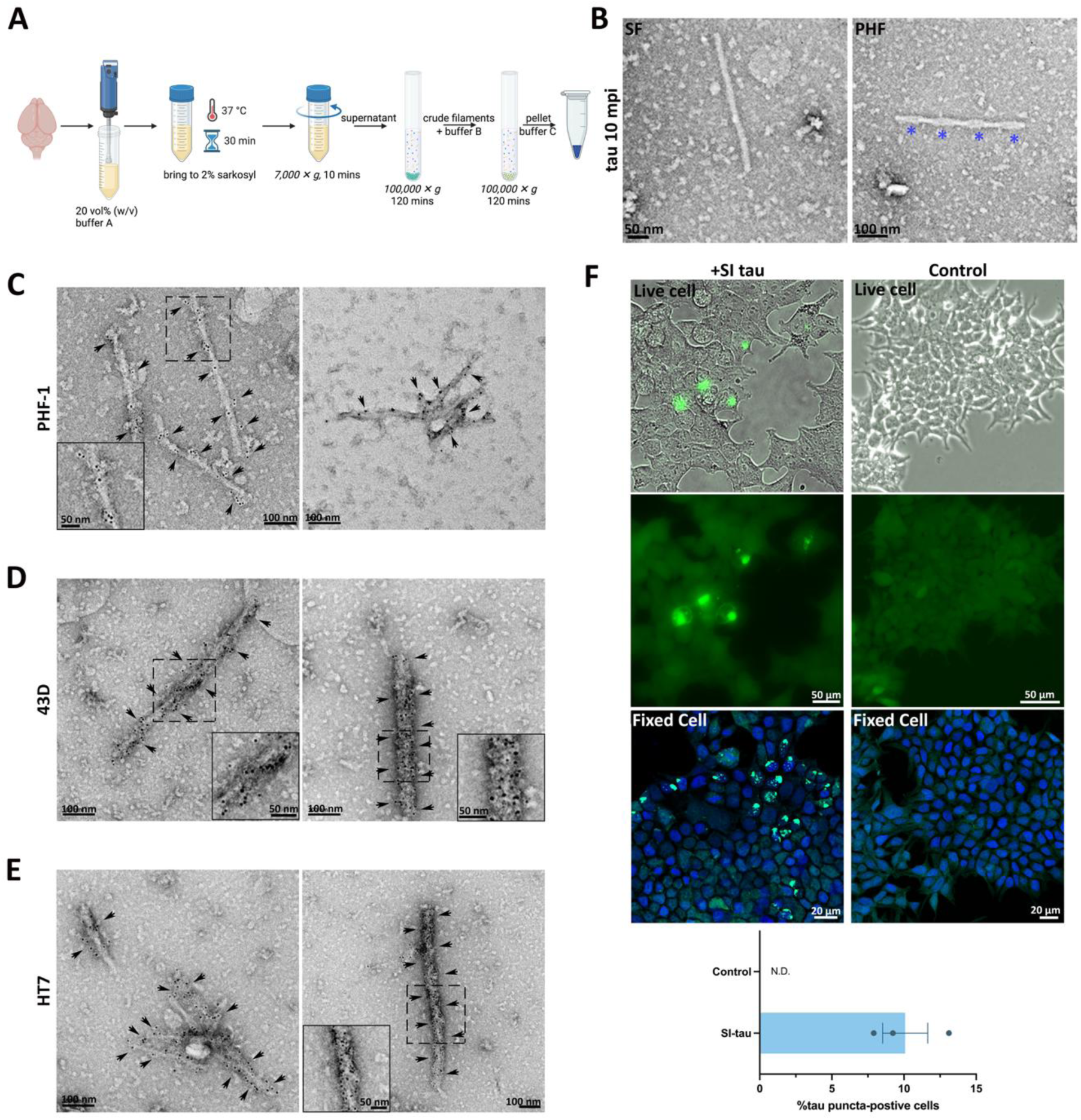
Tau pathology leads to neurodegeneration and glial responses in chimeric brains. **(A)** Representative hMAP2 and Ku80 double staining in cerebral cortex. Control chimeras show intact dendritic architecture, whereas tau-treated chimeras exhibit reduced overall hMAP2⁺ area and dendritic fragmentation. **(B)** Quantification of hMAP2⁺ area in hippocampus and cerebral cortex for control (n = 6), tau 6 mpi (n = 4), and tau 10 mpi (n = 8) chimeras. Data are mean ± SEM; Welch’s ANOVA with Dunnett’s T3 multiple-comparisons test; *p < 0.05, ***p < 0.001. **(C)** Representative hMAP2 and Homer1 double staining in hippocampus and cerebral cortex in control, tau 6 mpi, and tau 10 mpi chimeras. Homer1 puncta co-localized with hMAP2 are defined as postsynaptic structures associated with human neurons. **(D)** Quantification of Homer1 puncta co-localized with hMAP2 in hippocampus and cerebral cortex. Control, n = 6; tau 6 mpi, n = 4; tau 10 mpi, n = 4. Data are mean ± SEM; one-way ANOVA with Tukey’s multiple-comparisons test; *p < 0.05, ****p < 0.0001. **(E)** Representative GFP, NeuN, and AT8 triple labeling in cortex from control and tau 10 mpi chimeras. **(F)** Quantification of human neuron density within AT8⁺ regions (human neurons per AT8⁺ area), measured in the cortex and hippocampal formation. Control, n = 6; tau 6 mpi, n = 3; tau 10 mpi, n = 4. Data are mean ± SEM; one-way ANOVA with Tukey’s multiple-comparisons test; **p < 0.01, ****p < 0.0001. **(G)** Representative AT8 and phosphorylated MLKL (pMLKL) double staining in control and tau 6 mpi chimeras. Arrows indicate pMLKL⁺ neurons. **(H)** Representative AT8 and p62 double staining in control and tau 10 mpi chimeras. p62 accumulation in tangle-bearing neurons. **(I-K)** Representative of AT8 and Iba1 double labeling from control chimera and tau 10 mpi chimera. **(I)** Microglia were found surrounding the punctate AT8-positive phospho-tau deposits. Asterisks point out microglia phagocytosing tau. **(J)** Arrows highlight microglia-associated AT8^+^ tau puncta, indicating microglia actively engulfing phospho-tau. **(K)** Imaris-based 3D reconstruction further indicates microglial engulfment of AT8⁺ deposits. **(L)** Representative AT8 and GFAP double staining in control and tau 10 mpi chimeras. Asterisks denote reactive astrocytes associated with punctate AT8⁺ deposits; the inset indicates astrocytic processes surrounding an AT8⁺ tangle. Scale bars, as indicated in the panels.

Human neuron loss was further assessed by quantifying NeuN-positive neurons in the hippocampus and cortex, using GFP or Ku80 to identify human cells (**Fig. 4E**, Supplementary **Fig. S4A**). We observed a significant reduction in NeuN^+^ human cell number in tau-treated animals compared with controls (Control: 1712±72.20; tau_6mpi: 1201±17.97; tau_10mpi: 523.2±108.4) (**Fig. 4F**), indicating tau-associated human neuron loss.

To explore mechanisms underlying neuronal death, we examined markers of major cell death pathways and cellular homeostasis. Classic cell death markers, including cleaved Caspase-3 (apoptosis^48^), 4-HNE (ferroptosis^49^), and cleaved GSDMD (pyroptosis^50^), showed no significant differences between control and tau-treated chimeras (Supplementary **Fig. S4B-D**). The necroptosis marker pMLKL was detected in a subset of AT8-positive, tangle-bearing neurons (**Fig. 4G**), consistent with previous reports implicating necroptosis in tau-mediated neuronal loss^23^. However, the pMLKL signal was relatively sparse, suggesting that additional cell death mechanisms may also contribute. We further observed significant accumulation of the autophagy marker p62 in tangle-bearing neurons (**Fig. 4H**), consistent with previous research highlighting impaired autophagic flux and consequent metabolic stress accompanying tau pathology^51^, suggesting restoration of autophagy as a potential therapeutic strategy.

Finally, we evaluated glial response. Microglia formed barrier-like structures around neuropil threads and actively engulfed AT8-positive puncta (**Fig. 4I-K**). Additionally, astrocytes were closely adjacent to tangles (**Fig. 4L**). Together, these findings demonstrate that tau pathology drives progressive neurodegeneration and glial response in chimeric brains.

### Tau burden is reflected by plasma biomarkers and correlates with behavioral performance in chimeras

To evaluate whether the model recapitulates clinically relevant manifestations and provides translational value, we assessed plasma tau biomarker expression and cognitive function in chimeras. Plasma pT217-tau, an established clinical biomarker for AD-related tau pathology, was significantly elevated in tau seed–treated chimeras compared with controls at 6 mpi, with a modest further increase by 10 mpi (Control: 2.813±0.139; tau_6mpi: 3.498±0.238; tau_10mpi: 3.927±0.078) (**Fig. 5A**). These longitudinal trajectories mirror those reported in AD patients, in which pT217-tau level increases most sharply during early-to-mid stages and reaches a plateau at later stages^52^.

**Figure 5.**
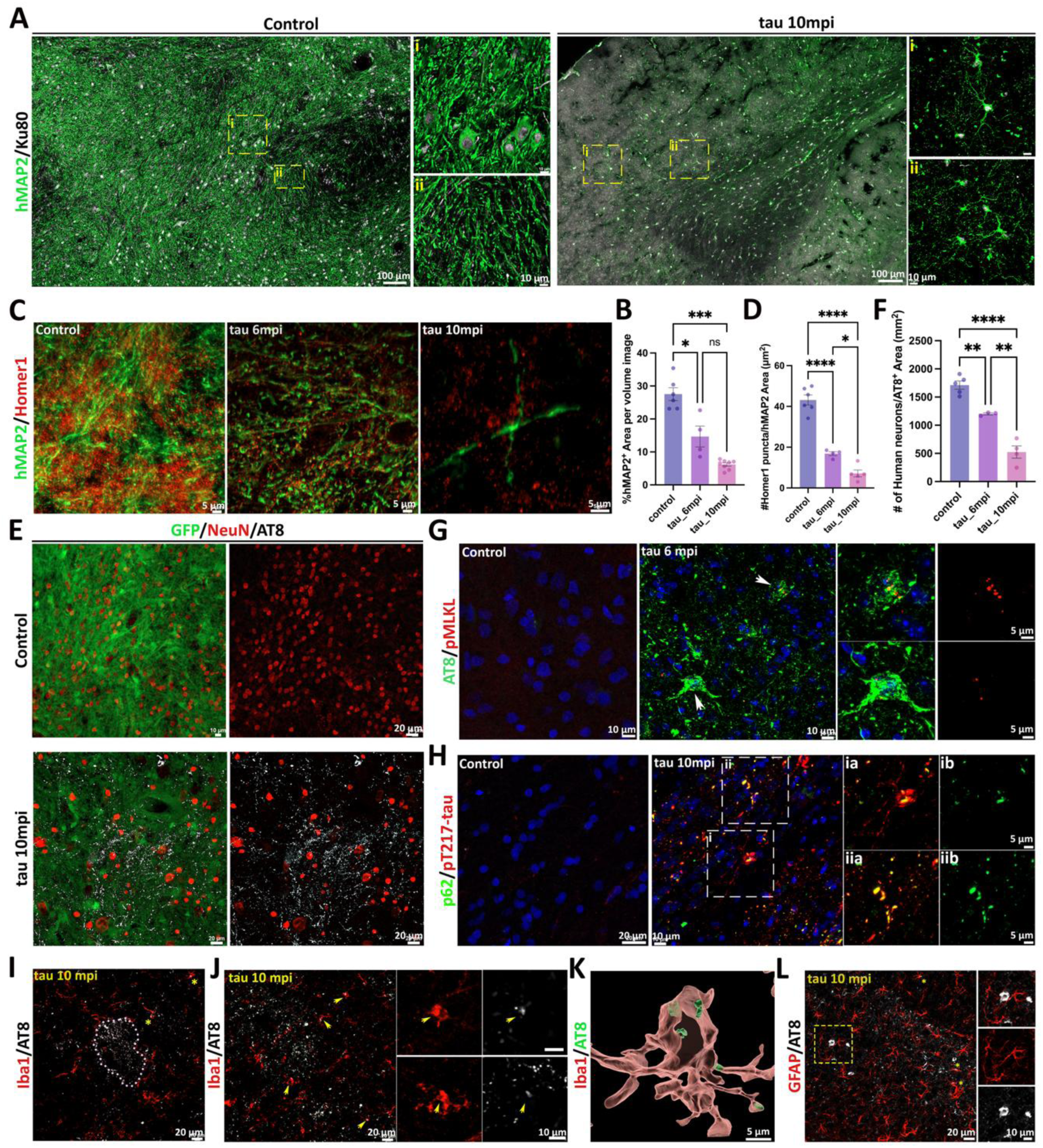
Progressive tau pathology is reflected by elevated plasma pTau-217 and correlates with behavioral performance in chimeras. **(A)** Plasma pTau-217 levels in 14-month-old control (n = 6), tau 6 mpi (n = 6), and tau 10 mpi (n = 6) chimeras. Data are mean ± SEM; One-way ANOVA with Tukey’s multiple comparison tests; ns, not significant, *p < 0.05, ***p < 0.001. **(B)** Open-field test schematic (left) and quantification of total distance traveled, and time spent in the center. N= 8- 20 mice in each group. Data are mean ± SEM; unpaired two-tailed t test between age-matched control and tau chimeras; ns, not significant. **(C)** Novel object recognition schematic (left) and quantification of total distance traveled and novel-object preference. N= 8- 20 mice in each group. Data are mean ± SEM; unpaired two-tailed t test between age-matched control and tau chimeras; ns, not significant; **p < 0.01. **(D)** Y-maze schematic (left) and quantification of total arm entries and spontaneous alternation. N= 8- 20 mice in each group. Data are mean ± SEM; unpaired two-tailed t test between age-matched control and tau chimeras; ns, not significant; *p < 0.05.

We next performed a panel of behavioral tests to assess cognitive function in chimeras at 4, 6, and 10 mpi. In the open field test, total travel distance and time spent in the center zone were similar between the groups of mice, indicating no deficits in locomotor activity or anxiety-like behavior in tau-treated mice at any time point (**Fig. 5B**). In the novel object recognition task, performance was comparable between control and tau-treated chimeras at 4 and 6 mpi. However, by 10 mpi, tau-treated animals showed a significant reduction in preference for the novel object, indicating impaired recognition memory after prolonged tau exposure (control vs. tau: 4mpi: 49.353±2.848 vs. 53.350±4.276; 6mpi: 51.668±5.863 vs: 48.792±5.564; 10 mpi: 50.556±3.484 vs. 37.245±2.812) (**Fig. 5C**). Consistently, in the Y-maze, 10 mpi tau-treated chimeras displayed a significant decrease in spontaneous alternation, suggesting deficits in short-term spatial working memory (control vs. tau: 4mpi: 48.484 ± 1.671 vs. 50.071 ± 2.148; 6mpi: 48.115 ± 3.124 vs. 43.551 ± 2.597; 10 mpi: 49.923±3.168, vs. 40.225±2.509) (**Fig. 5D**). Together, these findings demonstrate neuronal chimeras recapitulate clinically relevant features of AD tau pathology, including plasma pT217-tau and progressive cognitive decline, underscoring the translational utility of this model.

### Single-nucleus RNA sequencing reveals species-specific transcriptomic responses to tau pathology

Our results highlighted that human neurons are more vulnerable to tau pathology than neighboring mouse neurons (**Fig. 2**). To define human-specific transcriptional responses to tau pathology, we performed snRNA-seq on the whole brain, excluding cerebellum, from control and tau-seeded chimeras (Supplementary **Fig. S5A**). We identified 8 human-derived clusters and 9 mouse host clusters (**Fig. 6A**, Supplementary **Fig. S5D, E**). Human nuclei were annotated as excitatory neurons, inhibitory neurons, immature neurons, astrocytes, radial glia, oligodendrocytes, committed oligodendrocyte precursors (COP), and oligodendrocyte progenitor cells (OPCs) (**Fig. 6B, C**).

**Figure 6.**
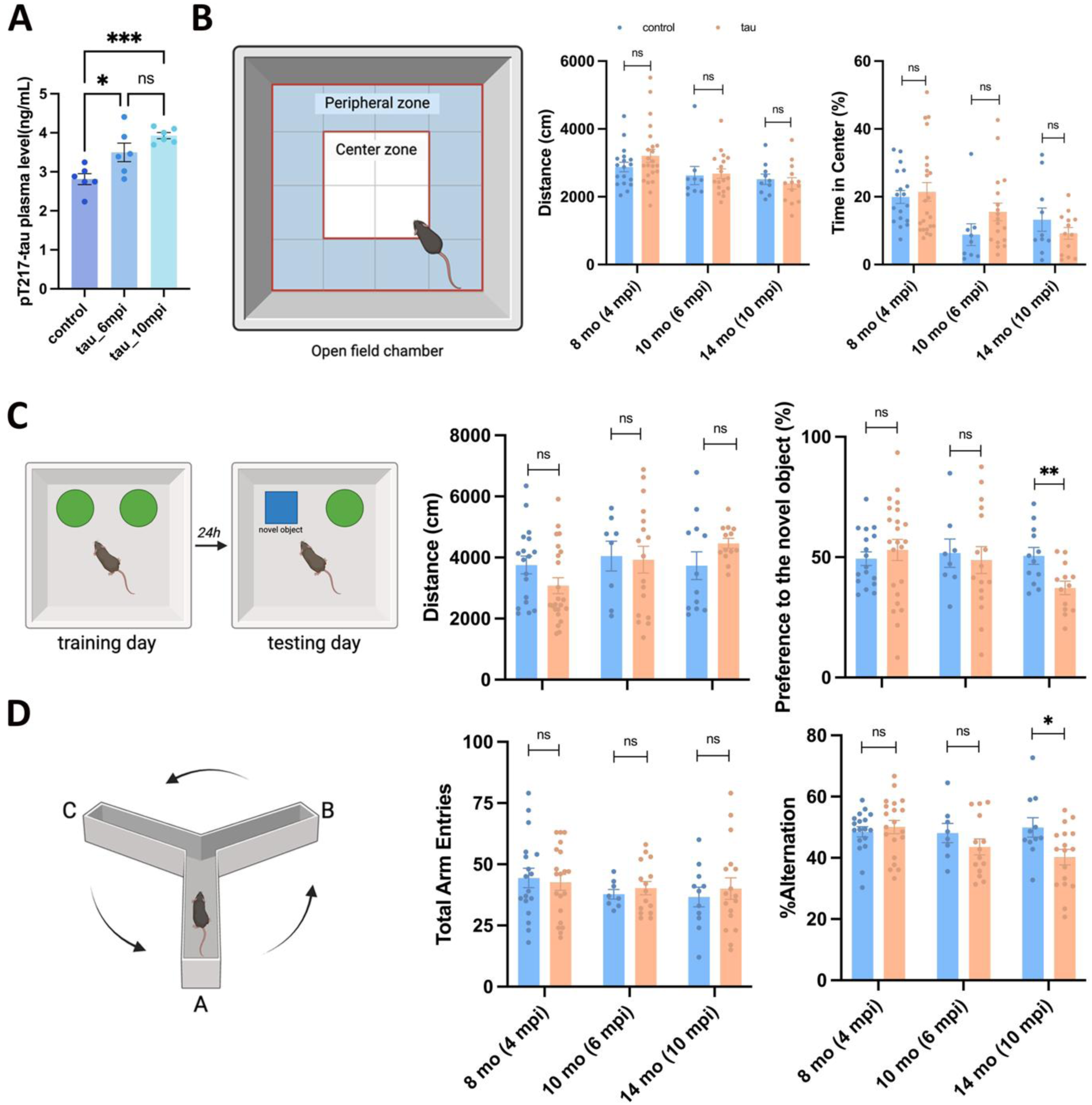
snRNA-seq reveals species-specific transcriptomic responses to tau pathology. **(A)** UMAP embedding of all nuclei profiled from chimeric brains after alignment to a combined human–mouse reference. Nuclei were assigned as human or mouse using a stringent species-mapping threshold (≥95% of reads mapped to the corresponding genome); details of quality-control filtering are in Supplementary **Fig. S5**. **(B)** UMAP of the human (transplanted) nuclei subset, colored by annotated cell classes. **(C)** Dot plot of canonical markers used for cell-type annotation. Dot size indicates the fraction of nuclei expressing each gene, and dot color indicates average expression (scaled within gene). **(D)** Volcano plot of differential expression in human excitatory neurons (tau vs control). Differentially expressed genes (DEGs) were defined as |log2FC| > 0.25 and adjusted p < 0.05. Representative genes are labeled and color-coded by putative functional categories. **(E)** Pathway enrichment analysis for significantly downregulated (left) and upregulated (right) genes in human excitatory neurons under tau pathology. The top 10 terms (adjusted p < 0.05) are shown; bar length denotes −log10(FDR), bars are shaded by overlap gene count, and the overlap ratio (overlap genes/genes in pathway) is indicated for each term. **(F)** Cross-species comparison of tau responses in excitatory neurons. A four-quadrant plot contrasts human and mouse log2FC (tau vs control), highlighting concordant and discordant gene regulation across species. Reactome pathway enrichment analysis was performed on genes in discordant quadrants (highly expressed in human or mouse neurons, respectively), and the top 15 significant terms (adjusted p < 0.05) are displayed. **(G)** Overlap tau-associated DEGs in transplanted human excitatory neurons with previously published AD patient excitatory neurons. Overlap proportions are shown for up- and downregulated gene sets (counts indicated). Reactome pathway enrichment analysis was performed on overlapping genes, and the top 15 significant pathways (adjusted p < 0.05) are shown along with a gene–pathway tile map indicating which overlapping genes contribute to each term. Published data were from Mathys et al. 2023 and Otereo et al.2022.

Since prior studies have shown that excitatory neurons are vulnerable to tau pathology^53–55^, we performed differential gene expression analysis in human excitatory neurons from tau-exposed versus control chimeras (|log2FC| > 0.25, adjusted *p* < 0.05) (**Fig. 6D**, Supplementary **Table S4**). Downregulated genes were centered on synaptic structure and neurotransmission, including presynaptic release machinery (*SNAP25, VAMP2, SYP*) and postsynaptic glutamatergic components (*GRIA1, GRIN2B*). *BDNF* was also significantly decreased, aligning with the reported attenuation of neurotrophic support in AD^56, 57^. In parallel, reduced *MAP2* and *HOMER1* reinforced our prior observation of dendrite and synapse loss (**Fig. 4A-D**). Upregulated genes were enriched for mitochondrial function (*MT-CO1, MT-ND1*), endosomal-lysosomal trafficking and lipid handling (*SORL1, ABCA1*), and calcium/cAMP signaling (*CAMK4, PPP3CA*), suggesting compensatory stress adaptation (**Fig. 6D**, supplementary **Table S4**). Reactome pathway analysis confirmed this pattern: downregulated pathways were signaling-centered (transmission across chemical synapses, neurotransmitter receptors and postsynaptic signal transmission), while upregulated pathways were dominated by mitochondrial RNA processing and stress remodeling (FASTK family–associated pathways regulating mitochondrial RNA processing/stability, mitochondrial RNA degradation) (**Fig. 6E**, supplementary **Table S5**).

To identify species-specific transcriptomic responses to tau pathology, we mapped human gene symbols to one-to-one mouse orthologs and computed a difference-in-differences metric (ΔΔlog2FC) per gene, capturing tau-induced divergence or convergence between human and mouse neurons beyond intrinsic species differences (Supplementary **Table S6**). The resulting concordant and discordant responses were visualized in quadrant plots comparing tau-versus-control log2 fold changes in human versus mouse neurons (**Fig. 6F**). Concordant genes revealed a shared core tau response between human and mouse neurons, though synaptic genes such as *BDNF* and *HOMER1* were more strongly suppressed in human neurons. Among discordant genes, human-up/mouse-down genes were enriched for RNA metabolism, splicing, and proteostasis pathways, whereas human-down/mouse-up genes were enriched for neuronal/synaptic modules (chemical synaptic transmission, receptor/postsynaptic signaling, axon guidance, and Rho GTPase pathways). These results suggest that under the same tau challenge, mouse neurons relatively preserve synaptic programs that are suppressed in human neurons, while human neurons exhibit stronger stress response.

Finally, we benchmarked our chimera model against patient-relevant transcriptional signatures by comparing DEGs from tau-exposed versus control human excitatory neurons with those from published AD human excitatory neuron datasets^58, 59^. The analysis demonstrated robust directional concordance, with approximately one-third of DEGs overlapping (Exc_Up: 214/581; Exc_Down: 395/1121) (**Fig. 6H**, Supplementary **Table S7, S8**). Shared downregulated genes converged on synaptic transmission across chemical synapses and glutamatergic signaling (*SNAP25, STXBP1, GRIA1, GRIN2B, NRXN1/NRXN3, SHANK3*), while shared upregulated genes converged on mitochondrial RNA-processing and AD-relevant stress responses (*PTK2B, ABCA1, RYR2, CAMK4*). This cross-dataset concordance supports the translational relevance of the chimera model, which recapitulates a conserved AD-like program in human neurons.

### PSEN2 N141I mutation exacerbates tau pathology and synaptic loss in chimeras

To explore the utility of neuronal chimeras for disease modeling, we investigate how a genetic risk factor influences tau pathology. We focused on *PSEN2* N141I, a rare cause of familial AD known to alter γ-secretase processing and increase Aβ production^60^. Given that amyloid pathology is thought to synergistically promote tau pathology^61^, we sought to determine whether this mutation can trigger the formation of Aβ aggregates and accelerate tau pathology *in vivo*.

NPCs carrying the *PSEN2* N141I mutation or an isogenic control were transplanted into host brains and exposed to an identical dose of P-tau seeds. AT8 staining revealed more extensive tau pathology in *PSEN2* N141I chimeras across hippocampus, cerebral cortex, and entorhinal cortex at 10 mpi (**Fig. 7A**). Quantification confirmed a significant increase in AT8⁺ neurons relative to isogenic controls (isogenic control: 2.855 ± 0.306; *PSEN2* N141I: 4.640±0.364) (**Fig. 7B**). Co-staining with Ku80 or 43D verified that the majority of tangles arose from human neurons (**Fig. 7C, D**).

**Figure 7.**
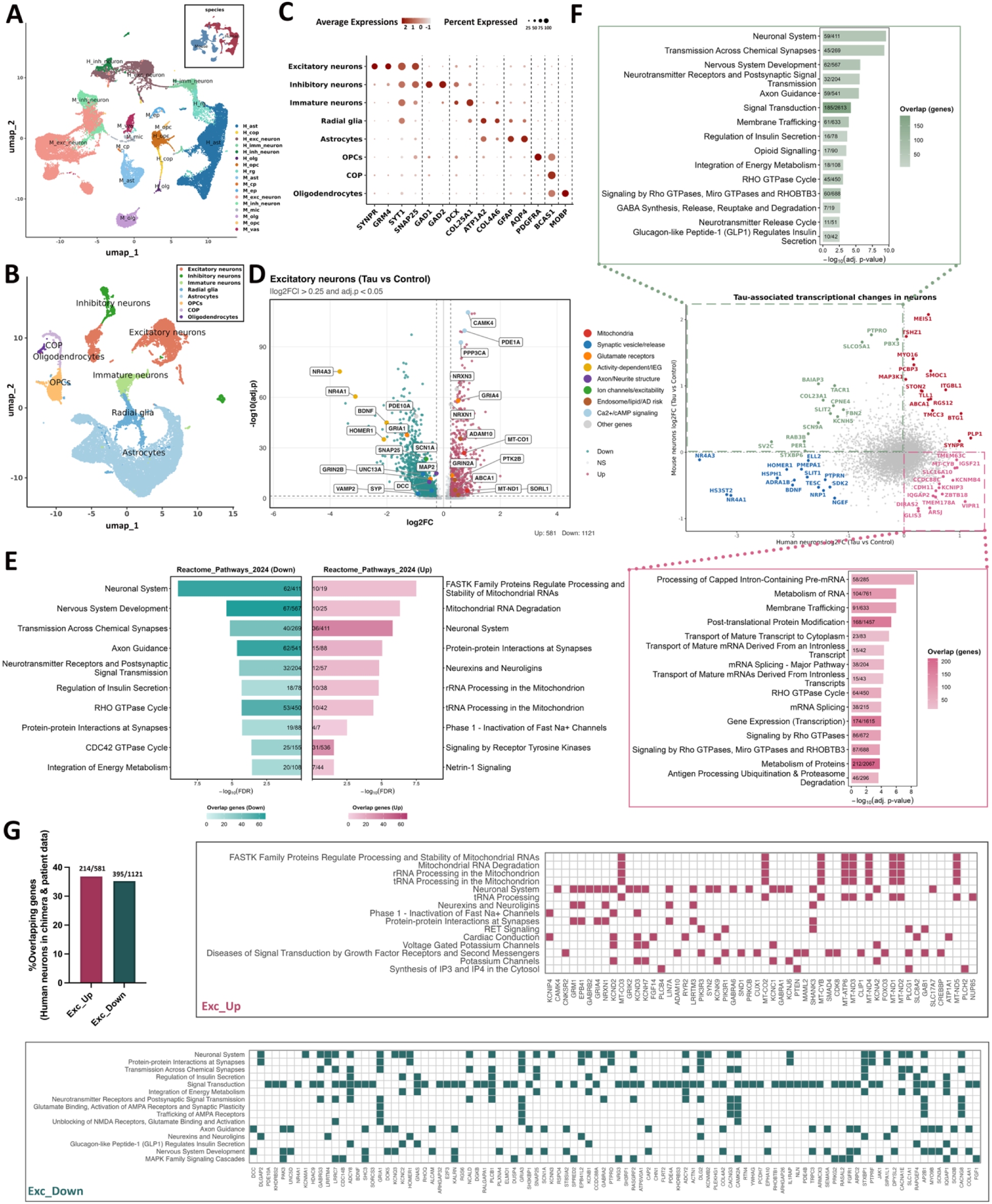
The PSEN2 N141I mutation exacerbates tau pathology and synaptic loss in chimeras. **(A)** Representative AT8 immunostaining at 10 months post tau-seed exposure in chimeras transplanted with NPCs from isogenic control or PSEN2 N141I mutant lines. Images are shown from entorhinal cortex (EC), hippocampus (HIP), and cortex (CTX). **(B)** Quantification of the number of AT8⁺ cells in isogenic control or PSEN141I groups. Data are mean ± SEM; unpaired two-tailed t test, *p < 0.05. **(C)** Representative of MC1 and Ku80 double staining in PSEN2 N141 chimeras at 10 months post tau injection. **(D)** Representative 43D, pTau-217, and Ku80 triple labeling in PSEN2 N141I chimeras at 10 months post-injection. Arrows mark 43D⁺ tangle-bearing human neurons; asterisks denote 43D and Ku80 positive human neurons without detectable tangles. **(E)** Representative MOAB2 and pTau-217 double staining in PSEN2 N141I chimeras at 10 months post-injection. Arrows indicate co-localization of intraneuronal Aβ (MOAB2) with hyperphosphorylated tau (pTau-217). **(F)** Representative hMAP2 and Homer1 double staining at 10 months post-injection in isogenic control versus PSEN2 N141I chimeras. The hMAP2⁺ area and the number of Homer1 puncta co-localized with hMAP2 were quantified. Data are mean ± SEM; unpaired two-tailed t test; ns, not significant; *p < 0.05. **(G)** Representative AT8, NeuN, and Ku80 triple staining at 10 months post-injection in isogenic control versus PSEN2 N141I chimeras. Human neuron density within AT8⁺ regions (Ku80⁺NeuN⁺ cells per AT8⁺ area) was quantified in the cerebral cortex and hippocampus. Data are mean ± SEM; unpaired two-tailed t test; ns, not significant. **(H)** Representative p62 and pTau-217 double staining at 10 months post-injection in isogenic control versus PSEN2 N141I chimeras. p62⁺ area per pTau-217⁺ tangle was quantified. Data are mean ± SEM; unpaired two-tailed t test; *p < 0.05. Scale bars, as indicated in the panels.

Despite the exacerbated tau pathology, extracellular Aβ plaques were not detected (Supplementary **Fig. S7**), and only sparse intraneuronal Aβ aggregates were observed adjacent to tau lesions in the hippocampus (**Fig. 7E**), suggesting that *PSEN2*-mediated augmentation of tau pathology is independent of extracellular Aβ deposition in this model. To assess associated neurodegeneration, we examined synaptic and dendritic integrity. Homer1^+^ puncta localized to hMAP2^+^ dendrites were significantly reduced in *PSEN2* N141I chimeras (isogenic control: 6.467±0.319; *PSEN2* N141I: 1.873±1.022) (**Fig. 7F**), whereas dendritic loss was comparable between groups (isogenic control: 7.253 ± 0.310; *PSEN2* N141I: 5.547±1.327) (**Fig. 7F**), likely because isogenic control already exhibited substantial dendritic fragmentation and loss under seeded tau pathology. Human neuron density within AT8⁺ regions showed a downward trend in *PSEN2* N141I chimeras that did not reach statistical significance (isogenic control: 615.3±80.96; *PSEN2* N141I: 539.9±85.93) (**Fig. 7G**). Finally, given that PSEN2 mutations have been linked to autophagosome–lysosome dysfunction^62^, we examined p62 accumulation and found an significantly greater p62-positive area per pT217-tau⁺ tangle in *PSEN2* N141I neurons (isogenic control: 12.17±0.798; *PSEN2* N141I: 18.70±1.398)(**Fig. 7H**), implicating that impaired autophagic clearance may be associated with the accelerated tau pathology. Together, these data indicate that *PSEN2* N141I amplifies tau pathology and synaptic loss in chimeras independent of extracellular Aβ aggregates.

## Discussion

The development of tau-targeted therapies in AD has been impeded by current models that inadequately replicate authentic disease phenotypes, particularly the mature forms of AD tau pathology, and by a limited understanding of species-specific responses to tau pathology^4, 63, 64^. Here, we develop a patient-derived tau-seeded human neuronal chimera model that faithfully recapitulates adult tau isoform composition and mature AD tau pathology, and reveals human-specific vulnerability to tau pathology. In this system, human neurons mature *in vivo*, expressing all six adult tau isoforms at a balanced 3R:4R ratio, and develop robust, mature AD tau pathology, accompanied by neurodegeneration and cognitive decline. Importantly, this model uncovers a striking human-specific neuronal vulnerability to tau pathology, linked to distinct transcriptional responses in human neurons associated with tau uptake and synaptic dysfunction. Together, this work establishes a human-relevant *in vivo* platform that enables direct interrogation of species-specific disease mechanisms and advances translational therapeutic development. By revealing intrinsic differences in neuronal vulnerability and response to tau pathology, our study offers new insights into translational challenges and highlights human-specific pathways as promising targets for therapeutic intervention.

Compared with existing models, the patient-derived tau-seeded human neuronal chimera model provides a more physiologically relevant platform to study tauopathies, particularly AD. First, these chimeras enable human neurons to mature and recapitulate adult human tau features, including all six tau isoforms with an approximately 1:1 3R: 4R ratio, as observed in the adult human brain. This is critical, as both isoforms contribute to AD tau pathology, whereas adult rodents predominantly express 4R tau^16, 17,19^. Human *MAPT* knock-in mice provide powerful tools to recapitulate human tau isoforms, yet they are constrained by species-specific differences in neuronal biology^65, 66^. In contrast, within this long-term, physiologically relevant environment, human neurons progressively mature in chimeric brains, exhibiting spontaneous action potentials, functional integration into host neural circuits^28^, and aging-associated features, including senescence markers (Supplementary **Fig. S1K**). Consistently, chimeric brains show an age-dependent increase in 4R tau expression (**Fig. 1E, F**) and achieve an adult-like 3R:4R ratio with all six tau isoforms by 6 months of age (**Fig. 1G**).

Second, by using patient-derived tau seeds rather than FTD-associated *MAPT* mutations, chimeric brains accurately model sporadic tauopathies and uniquely recapitulate mature AD tau pathology, a key feature lacking in existing models. Although pathogenic *MAPT* mutations drive tau pathology, they do not reflect etiological drivers of tau pathology in AD and are associated with distinct clinical, pathological, and behavioral phenotypes^67, 68^. Notably, tau aggregates formed in *MAPT* mutation–based models differ structurally from those observed in human AD brains^69^. Prior studies show that tau seeds induce tau pathology in animals^32, 70, 71^. Here, we demonstrated that patient-derived tau seeds can robustly induce mature AD tau pathology in human neurons in chimeras. Although oligomeric tau species may initiate toxicity, they are transient and difficult to define^72, 73^. In contrast, mature tau pathology is strongly associated with neurodegeneration, cognitive decline, and disease severity, and provides a stable and quantifiable readout of disease progression and therapeutic response^9, 45, 74^. To date, existing models are more accurate when dissecting early stage AD tau pathology, while recapitulation of mature tau pathology, characteristics of advanced-stage human AD, remains as a limitation. For example, microRNA-reprogrammed neurons^24, 25^ and neuronal chimera models with human neurons transplanted into amyloid-rich environments^22, 23^ capture key aspects of human tau biology but typically generate limited or incomplete NFTs containing only SFs. In contrast, the patient tau-derived neuronal chimeras faithfully recapitulate mature AD tau pathology, including NFTs and neuropil threads composed of PHFs and SFs containing 3R and 4R tau. Furthermore, the pathology accumulates and spreads across anatomically connected regions, accompanied by neurodegeneration, elevated plasma tau biomarker, and memory deficits. (**Fig. 3I**, **Fig. 4, 5**). Together, these features establish neuronal chimeras as a physiologically relevant and powerful platform for modeling AD tau pathology and therapeutic testing. Beyond AD tau pathology, this chimera seeding model can be extended to study other tauopathies and neurodegenerative proteinopathies by incorporating disease-relevant seeds^75^, such as Parkinson’s disease (α-synuclein)^76^, amyotrophic lateral sclerosis (ALS) and FTD (TDP-43)^77^, Huntington’s disease (hungtingtin)^78^, as well as amyloid-β in AD^79^.

A key finding of this study is the intrinsic vulnerability of human neurons to tau pathology. The growing failure of tau-targeted therapies that showed promise in animal models highlights the need to define human-specific disease mechanisms to guide more effective therapeutic development^4, 5^. In chimeric brains where human and mouse neurons coexist under identical conditions, tau pathology arose predominantly in human neurons (∼75%), indicating the same seed dose that robustly induced pathology in human neurons was suboptimal in mouse neurons, consistent with prior observations that wild-type mice typically require larger tau seed quantities and longer time courses^32^. Supporting this, when identical seeds were injected into *Rag1^⁻/⁻^* mice lacking human cells (2 μg), only minimal AT8 staining was observed at 10 mpi (Supplementary **Fig. S2G**). Mechanically, snRNA-seq revealed that several tau uptake receptors, including LRP1, HS3ST1, and SDC3, are more highly expressed in human excitatory neurons under baseline conditions (**Fig. 2E**), and *in vitro* tau endocytosis assays confirmed significantly greater uptake in human versus mouse neurons (**Fig. 2F, G**). Cross-species transcriptomic comparison further showed that tau exposure produced more pronounced synaptic program suppression and RNA/proteostasis remodeling in human neurons, while mouse neurons showed relative preservation of synaptic and axonal programs (**Fig. 6F**). Together, these findings indicate that human neuronal vulnerability reflects both greater tau uptake capacity and heightened sensitivity to tau-driven cellular disruption, underscoring the importance of human models for defining human-specific disease mechanisms and informing therapeutic strategies that target human-specific pathways.

To explore the potential of neuronal chimeras for disease modeling, we introduced the PSEN2 N141I mutation into transplanted NPCs to test whether endogenous Aβ production could modulate seeded tau pathology. PSEN2 was chosen over PSEN1 because, unlike the highly penetrant and clinically aggressive PSEN1 mutations that often present with atypical features^80–82^, PSEN2 mutations more closely recapitulate typical AD pathology with later onset and lower penetrance^83^. This makes PSEN2 N141I particularly well-suited for studying Aβ–tau interactions in a model designed to capture core features of AD. By 10 months after tau inoculation, PSEN2 N141I chimeras showed exacerbated tau pathology compared with their isogenic controls (**Fig. 7A, B**). Despite this, we did not detect extracellular Aβ deposits (Supplementary **Fig. S7**) and observed only rare intracellular Aβ aggregates near tau lesions (**Fig. 7E**), which may reflect the milder nature and later onset of PSEN2-linked disease. We also observed minimal spatial overlap between intraneuronal Aβ and tau aggregates, suggesting that, in this context, extracellular Aβ aggregates are not the primary drivers of the more severe pathology. Instead, we found increased p62 accumulation in PSEN2 N141I neurons compared with isogenic controls (**Fig. 7H**), consistent with prior reports that PSEN2 mutations impair autophagosome clearance^62^.

The current patient-derived tau-seed neuronal chimera model also has limitations. Although tau pathology arises predominantly in human neurons, a subset of mouse neurons also develops pathology. To minimize potential contributions of murine tau to tau propagation, a useful future approach would be to transplant human neurons into MAPT knockout immunodeficient mice, in which tau pathology would arise exclusively from human neurons. Additionally, multiple factors contribute to tau pathology, including both neuron-autonomous factors^34, 84, 85^ and non-autonomous factors, such as microglia and adaptive immune cells (e.g., T cells)^86, 87^. The chimeric brain model enables targeted genetic manipulation of human neurons prior to transplantation (such as the PSEN2 study in **Fig. 7**) and is established in immunodeficient mice that retain innate immune cells (including microglia) but lack functional adaptive immune cells (T and B cells). This design can exclude the effects of adaptive immune cells and is particularly valuable for gaining new insights into neuron-autonomous mechanisms of tau pathology. However, understanding the contribution of adaptive immune cells will require future humanized models that incorporate these components. Overall, this patient-derived tau-seed neuronal chimera model recapitulates adult human tau features, reproduces mature AD tau pathology, and reveals species-specific vulnerability to tau pathology, establishing a valuable platform for species-specific mechanistic studies and therapeutic development.

## Materials and Methods

### Animals

*Rag1^-/-^* immunodeficient mice (B6.129S7-*Rag1*^tm1Mom^/J on a C57BL/6 background, the Jackson Laboratory) were used in this study. The mice were housed under a 12-h light/dark cycle, with access to food and water. All animal handling and use were conducted in accordance with the protocol approved by the Institutional Animal Care and Use Committee at Purdue University.

### Culture and quality control of hPSC lines and differentiation

Five different hiPSC cell lines, KOLF2.1, ND2.0, C5 control, UTY1, and PSEN2 N141I, including one hESC cell line CAGG, were used in this study (**Supplementary table S1**). The hPSCs were maintained under feeder-free conditions on hESC-qualified Matrigel (Corning) coated dish in mTeSR plus medium (STEMCELL Technologies). The hPSCs were passaged at approximately 70% confluency using ReLeSR (STEMCELL Technologies). The differentiation of hPSCs toward neural progenitor cells was performed as previously described^28, 88^. Briefly, dual inhibition of SMAD signaling was implemented to induce neural differentiation in embryoid bodies (EBs). EBs were cultured in neural induction medium composed of DMEM/F12 (HyClone) and 1 × N2 Supplement (Thermo Fisher Scientific) supplied with inhibitor SB431542 (2 μM, Stemgent) and noggin (50 ng/mL, Peprotech) for 7 days before plating on growth factor-reduced Matrigel (BD Biosciences) coated plates. EBs were cultured with neural induction medium supplied with laminin (1 μg/mL, Corning) for 7 days, and then NPCs in the form of neural rosettes were manually isolated. Isolated NPCs were expanded in NPC medium, which is composed of a 1:1 mixture of Neurobasal (Thermo Fisher Scientific) and DMEM/F12, supplemented with 1×N2, 1×B27-RA (Thermo Fisher Scientific), FGF2 (20 ng/mL, Peprotech), human leukemia inhibitory factor (hLIF, 10 ng/mL, Millipore), CHIR99021 (3 μM, Biogems), and SB431542 (2 μM). For *in vitro* tau uptake assay, NPCs were seeded on Poly-L-Ornithine (Sigma, P4957) and Laminin coated plates for further differentiation according to STEMCELL Technologies method with some minor modifications. Briefly, the cells were maintained with ND medium, which is composed of a 1:1 mixture of Neurobasal and DMEM/F12, supplemented with 1×N2, 1×B27-RA, BDNF (10 ng/mL, Peprotech), GDNF (10 ng/mL, Peprotech), L-Ascrobic acid (200 nM, Sigma Aldrich), and c-AMP (1 μM, Sigma Aldrich).

### Mouse embryonic stem cell culture and differentiation

Mouse embryonic stem cells were maintained following the manufacturer’s instruction. Briefly, irradiated MEF feeder cells were plated one day before thawing and plating of mouse ES cells. After the plating of mouse ES cells, cultures were maintained with KonckOut™ DMEM (Gibco, 10829018) supplemented with 15% KonckOut™ Serum (Gibco, 10828028) and mouse Leukemia Inhibitory factor (mLIF, Peprotech, 10 μg/mL), GluataMax™-I, 100X (1X, Gibco), 2-mercaptoethanol 1000X (1X, Gibco), and MEM™ Non-Essential™ Amino Acids Solution 100X (1X, Gibco). Neuron differentiation was performed in a similar approach as described above for hiPSCs, with mLIF substituting hLIF.

### Purification of Recombinant K18 Tau

Recombinant K18 P301L tau was purified as previously described for human WT 2N4R tau^89^. Briefly, E. coli BL21(DE3) cells were transformed with pT7-7 -K18 Tau, a gift from Dr. David Eliezer (Weill Cornell Medical College). Overnight cultures were used to inoculate 1L cultures supplemented with 100µg/mL ampicillin. Once an OD of 0.5 was reached, IPTG (1mM) was used to induce K18 tau expression for 4 hours. The cells were pelleted by centrifugation at 6,000g for 15 min at 4 °C, resuspended in lysis buffer (20 mM MES, 0.2 mM MgCl2, 1 mM EGTA, 0.1mM PMSF, 0.25 mg/mL lysozyme, 1 μg/mL DNase I, pH 6.8), and lysed with a French press cell disruptor at 4 °C. After boiling the lysate for 20 min, denatured proteins were pelleted by centrifugation at 30,000g for 30 min at 4 °C, and the supernatant was loaded onto a HiPrep SP HP column on an AKTA pure chromatography system. The column was equilibrated with cation exchange buffer (20 mM MES, 50 mM NaCl, 1 mM MgCl2, 1 mM EGTA, 2 mM DTT, and 0.1 mM PMSF, pH 6.8), and proteins were eluted with a linear gradient ranging from 50 mM to 1 M NaCl. The resulting protein solution was purified further using a HiLoad 16/600 Superdex 200 pg column with 1.5 column volumes of gel-filtration buffer (10 mM phosphate buffer, 2.7 mM KCl, and 137 mM NaCl, 1mM freshly added DTT pH 7.4), and fractions containing tau (identified by SDS-PAGE with Coomassie blue staining) were pooled and stored at −80 °C.

### Generation of Recombinant K18 Tau PFFs

Monomeric tau aliquots were thawed on ice. Prior to fibrillizing monomeric tau, the sample was filtered through a 100kDa MWCO filter to ensure the monomeric identity of the starting sample. Recombinant tau monomer was fibrillized in the presence of a 1:1 molar ratio of heparin (∼267µM heparin) at a concentration of 4mg/mL tau at 37°C at 1000rpm for 6 days in an orbital shaker. After fibrillization, aggregated tau was concentrated using a 10 kDa MWCO to 2mg/mL and stored in 25µL aliquots at -80°C. Tau concentration was determined by bicinchoninic acid assay (BCA), and gel filtration buffer was used to dilute tau to the desired concentrations.

### Preparation of pHrodo-labeled K18 Tau Fibrils

K18 tau PFFs were labeled via reactive amine labeling with pHrodo Deep Red TFP Ester at a molar ratio of 2.5:1 pHrodo: K18 in the dark for 1 hour. The labeled PFFs were then cleaned with the use of a 7kDa MWCO Zeba desalting column. Briefly, the column was placed in a 2mL microcentrifuge tube and centrifuged at 2500g to remove the storage solution. The column was then transferred to a new tube, and the labeled PFF sample was loaded onto the center of the resin bed. The sample was centrifuged at 1500g for 2 minutes, and the desalted sample was collected from the top of the resin bed.

### Tau uptake assay

Neuronal medium was removed and wells were washed once with DPBS. Neurons were then incubated in fresh ND medium containing 50 nM pHrodo-labeled K18 tau PFFs for 24 h at 37°C. After incubation, cultures were washed with DPBS to remove unbound PFFs and immediately imaged live on an EVOS M5000 microscope. Same exposure and imaging settings were kept consistent across conditions. Uptake was quantified in Fiji as the percentage of pHrodo-positive area within the field of view.

### Cell transplantation

Transplantation of NPCs into the brains of p0/p1 pups was performed as previously described^28^. In brief, three to five days before transplantation, frozen NPCs were thawed and maintained in NPC medium. On the day of transplantation, cells were dissociated using TryPLE and counted by Trypan Blue. Finally, cells were suspended at a 100,000 per μL in DPBS (Cytiva). Cryoanesthesia was performed by placing mouse pups on ice for 4 minutes. Pups were placed on a chilled digital stereotaxic device (RWD) equipped with a neonatal mouse adapter (Stoelting). Cells were transplanted by inserting the Hamilton needles through the skull into the target sites (coordinates from bregma: anteroposterior, -2.0 mm; lateral, ±1.0 mm; dorsoventral depth, -1.5 or -1.0 mm). 0.5 μL of cells was injected into each site. After transplantation, pups were allowed to recover on a heating pad. Fully recovered pups were returned to the cage. The pups were weaned at 3 weeks. No tumor formation was observed by the endpoint of the experiment.

### Purification and characterization of patient-derived pathological tau

Alzheimer’s disease postmortem brain tissue (frontotemporal region)-derived pathological tau was used as seeds to induce tau pathology in chimeric brains. Purification and characterization of pathological tau were described elsewhere^90^. In brief, 10% brain homogenate in homogenization buffer (20 mM Tris-HCl, pH 8.0, 0.32 M sucrose, 10 mM β-mercaptoethanol, 5 mM MaSO_4_, 1 mM EDTA, 10 mM glycerophosphate, 1 mM Na_3_VO_4_, 50 mM NaF, 2 mM benzamidine, 1 mM 4-(2-aminoethyl) benzenesulfonyl fluoride hydrochloride (AEBSF), and 10 μg/mL each of aprotinin, leupeptin, and pepstatin) was centrifuged at 27,000 × *g* for 30 min. The supernatant was further centrifuged at 235,000 × g for 30 min, and the resulting pellet, i.e., pathological tau-enriched fraction (P-tau), was collected and washed twice with saline and then resuspended in saline. The concentration of P-tau was confirmed using immuno-dot blot against serial dilutions of recombinant monomeric Tau40 (Supplementary **Fig. S2A**), as previously described^90^. To determine the seeding activity of P-tau, we performed *in vitro* seeded tau aggregation assay, in which HEK-293T cells expressing HA-tau_151-391_ were treated with P-tau. Seeded HA-tau_151-391_ aggregates were yielded by ultracentrifugation and analyzed by Western blots (Supplementary **Fig. S2A**), as previously described^90^.

### Intracerebral injection of adult mice

Human NPCs engrafted mice were injected with either pathological tau (disease group) or saline (control group) at four months of age. Before surgery, mice were weighed and anesthetized using a ketamine/xylazine cocktail at 100/10 mg/kg. Surgeries were performed on a RWD stereotaxic apparatus. P-tau or saline was aseptically injected into hippocampus CA1 (coordinates from bregma: anteroposterior, -2.5 mm; lateral, ±2.0 mm; dorsoventral depth, -1.8 mm). 2 μL of P-tau or saline was injected at each site. After suturing, mice were subcutaneously injected with Buprenorphine ER (0.5 mg/kg) for analgesia. Mice were returned to the cage and placed on a heating pad set to body temperature. After injection, mice were monitored twice daily for 72 hours.

### Sample isolation

Animals were euthanized by CO_2_ asphyxiation. For immunohistochemistry, mice were transcardially perfused first with PBS followed by 4% PFA in PBS. Brains were collected and fixed overnight in 4% PFA. Gradient dehydration was performed by sequentially immersing brain samples into 20% and 30% sucrose. Brains were embedded in Optimal Cutting Temperature (OCT) Compound and cut into 30 μm thick sagittal serial sections using a Epredia™ CryoStar™ NX50 Cryostat. Sections were mounted directly onto slides and stored at -20 °C until use. For RNA extraction, nuclei suspension preparation, and tau fibrils isolation, animals were perfused with ice-cold DPBS. The brain samples were quickly isolated and rinsed in ice-cold DPBS. Then the samples were snap-frozen in liquid nitrogen and stored in -80 °C until use. Blood samples were collected in EDTA-coated tubes before perfusion. Whole blood samples were centrifuged at 2000 × *g*, and the plasma supernatant was collected, aliquoted, and stored in -80 °C until use.

### RNA extraction and semi-quantitative PCR

For 3R and 4R tau expression level analysis, frozen brain samples of xenografts at 3, 4, 5, and 6 months of age (n =3 for each group) were used for RNA extraction using TRIzol Reagent (Invitrogen). Complementary DNA was prepared with SuperScript IV First-Strand Synthesis System (Invitrogen). Semi-quantitative PCR was performed to assess the RNA expression of 3R and 4R tau isoforms using the primers as previously described^24^ and listed in **Supplementary Table S2**. Gel images were captured using Azure Biosystem C280. Acquired images were quantified for the ratio of 3R and 4R tau isoforms using Photoshop.

### Western Blot analysis

To analyze the expression pattern of human tau protein in xenografts, frozen brain samples were homogenized in T-PER tissue protein extraction reagent (ThermoScientific, 78510) supplied with protease inhibitor cocktail (Cell Signaling Technology, 5871). The samples were then centrifuged at 10,000 × *g* for 5 mins to pellet the tissue debris. The supernatant was collected, and BCA protein assay was performed to determine protein concentration. Samples were then normalized with T-PER/protease inhibitor and treated with FastAP™ Thermosensitive Alkaline Phosphotase (EF0651, 10U FastAP™ Thermosensitive Alkaline Phosphotase 1 hr, 37 °C) according to the manufacturer’s instructions. Human brain whole tissue lysate (Biotechne, NB820-59177) was used as a positive control, and non-engrafted mice were used as a negative control. 4 × Laemmli Sample Buffer (Bio-Rad, 1610747) with 5% β-mercaptoethanol was added and samples were heated to 95 °C for 10 mins then loaded onto a 7.5% Mini-PROTEAN^®^ TGX™ Precast Protein Gel (Bio-Rad), with Tau Peptide Ladder (rPeptide T-1007-2) and Spectra™ Multicolor Broad Range Protein Ladder (ThermoScientific, 26634). After running, the gel was transferred onto a PVDF membrane (Bio-Rad, 1620174) using Trans-Blot^®^ Turbo™. Membrane was blocked with 5% BSA in TBS/0.1% Tween20 for 1 hr at RT and then incubated with primary antibody 43D (provided by Dr. Liu, 1:500) overnight at 4 °C. The next day, the membrane was washed 3 times with TBST, then treated with peroxidase-conjugated donkey anti-mouse secondary antibody (Jackson ImmunoResearch, 715-005-151) in TBST for 1 hr at RT. The membrane was washed 3 times with TBST, developed with an ECL system (Thermo Scientific), and imaged on a Bio-Rad imager.

### Immunofluorescence and imaging

For immunofluorescence analysis, sections were blocked with blocking solution (5% goat or donkey serum in PBS with 0.8% Triton X-100) at room temperature (RT) for 1 hr. Tissues were incubated with primary antibodies were diluted in the same blocking solution at 4 °C overnight. The sections were then washed three times with PBS (10 min each) and incubated with secondary antibodies for 1 hr under RT. After washing with PBS for three times (10 min each), the slides were mounted with the anti-fade Fluoromount-G medium containing 1,40,6-diamidino-2-phenylindole dihydrochloride (DAPI) (Southern Biotechnology). The primary antibodies and secondary antibodies used were listed in **Supplementary Table S3**. Slides were kept at 4 °C until imaging. Confocal images were obtained using a Zeiss LSM900 Confocal Microscope. Images were obtained using a 10x (0.45 NA) or 20x (0.8 NA) objective lens, and images of interest were acquired using ZEN 3.1 software. Images were processed using Fiji/Image J software. For quantification, three to five sections of each brain samples were randomly selected and imaged. For anatomical representative images on the spread and progression of tau pathology, a score from 0 to 3 was added by visually assessing all the anatomical regions on the atlas. Visualization of the data was performed using BrainGlobe-Heatmap^91^. Data are pooled from chimeras transplanted with different cell lines.

### Gallyas Silver Staining

The Gallyas silver staining was performed as previously described to stain Tau tangles^92^. Briefly, slides were washed in distilled water for one minute and transferred immediately to 5% potassium permanganate for 15 minutes. Next, slides were washed for 1 minute in 2% oxalic acid, followed by three washes in distilled water for 1 minute. Then the slides were incubated in alkaline silver iodide solution for 2-5 minutes followed by three rinses in 0.5% acetic acid for 1 minute each. Then the slides were incubated in physical developer solution for 15 -20 minutes. The developer was made freshly before use in a 1:1 ratio (solution A: solution B; solution A: 0.2 g ammonium nitrate, 0.2 g silver nitrate, 1 g tungstosilicic acid, 0.5 mL 37% formaldehyde solution in 100 mL distilled water; solution B: 5 g Anhydrous sodium carbonate in 100 mL distilled water). Next, the samples were washed in 0.5% acetic acid for 3 times, 1 minute each. Subsequently, samples were washed in distilled water for five minutes, and incubated in 0.5% gold chloride solution, and sodium thiosulfate for 5 minutes each. Finally, the samples were washed in distilled water for five minutes, counterstained with nuclear fast red, and mounted using Permount solution (Fisher, SP15-100).

### Sarkosyl-insoluble tau extraction from P-tau injected chimeras

Sarkosyl-insoluble tau was extracted from chimeras 10 months post P-tau injection as described previously. Tissues were homogenized in 20 vol% (w/v) buffer A (10 mM Tris-HCl pH 7.4, 0.8 M NaCl, 10% Sucrose and 1 mM EGTA), brought to 2% sarkosyl and incubated at 37 °C for 30 min. The samples were then centrifuged at 7,000 × *g* for 10 mins, followed by spinning of the supernatant at 100,000 × *g* for 120 mins. The pellets were resuspended in buffer A and were diluted threefold in buffer B (50 mM Tris-HCl pH 7.5, 0.15 M NaCl, 10% sucrose and 0.2% sarkosyl), followed by a 120-min spin at 100,000 × *g.* The pellets were then resuspended in 100 μL g^-1^ buffer C (20 mM Tris-HCl pH 7.4 and 100 mM NaCl), aliquoted and snap-frozen in liquid nitrogen. The proteins were then stored in -80 °C until use.

### Negative staining and Immune-Electron Microscopy

Immunogold labeling and negative staining of sarkosyl insoluble tau extracted from chimeras were performed as described before^93, 94^. For immune-EM, 5 μL diluted sarkosyl-insoluble fractions of tau filaments were loaded onto glow-discharged 400 mesh carbon film-coated copper grids for 5 mins and washed three times with PBS, then blocked for 30 mins with 5% acetylated BSA (Aurion, 25552-02). The grids were incubated with primary antibodies diluted in incubation buffer (0.1% BSA-c Aurion, 1:20) at RT for 2 hours, followed by three washes in incubation buffer. Then the grids were incubated with 6 nm gold-conjugated secondary antibody (1:20, Aurion) in incubation buffer for 2 hrs at RT. Finally, grids were washed three times with deionized water and negatively stained with 2% uranyl acetate for 1 minute. The grids were dried and imaged using TECNAI G2 20 TEM, equipped with a bottom-mounted Gatan US1000 2Kx2K.

### *In vitro* seeding assay using chimeras-derived sarkosyl-insoluble tau

To assess the seeding capacity of the sarkosyl-insoluble tau extracted from P-tau injected chimeras, we performed *in vitro* seeding assay using HEK293 biosensor cells, a gift from Dr. Marc Diamond (UT Southwestern Medical Center). Briefly, the cells were thawed and maintained in HEK medium (DMEM high glucose (Corning), 10% FBS (Gibco), 1% Non-Essential Amino Acids (Gibco)). The day before the seeding experiment, cells were replated on a Poly-D Lysine-coated cover glass in a 24-well plate at a density of 20,000 cells/well and left to adhere overnight. The following day, cells were seeded with chimeras-derived sarkosyl-insoluble tau. The seeds were bath sonicated for 2 mins at 80% power before being diluted into Opti-MEM (Gibco). The seeding mix was prepared using Lipofectamine stem (Invitrogen) according to the manufacturer’s instructions, with 5 μL of SI-tau mixed with 1 μL of Lipofectamine stem reagent. For the control group, buffer C (employed to resuspend sarkosyl insoluble tau) was added to the seeding mix as a negative control. Each condition was done in triplicate. Cells were monitored daily and were fixed for immunofluorescence analysis at approximately 70% of confluency.

### Plasma pT217-tau measurements

The plasma level of phospho-tau 217 was measured using ELISA kit (Abcam, ab318936). Plasma samples were thawed at room temperature and diluted fourfold using the kit-provided sample diluent. The standards were prepared per the manufacturer’s instructions. Samples from control, P-tau 6 months post injection, and P-tau 10 months post-injection were analyzed (n = 6 for each group). All measurements were done in triplicate.

### Behavioral tests

To assess the progressive memory deficits in chimeras, we performed behavioral tests on control chimeras and P-tau injected chimeras at three different time points: 8 months old (4 months post P-tau injection), 11 months old (7 months post P-tau injection), and 14 months old (10 months post P-tau injection). Prior to the behavior tests, animals were handled daily for seven consecutive days to minimize possible anxiety caused by handling. The open field chamber and Y-maze were wiped with 70% ethanol and allowed to dry completely between animals to eliminate odor effects. All behavior tests were performed by an investigator who were blinded to treatments. SMART 3.0 (Panlab) software was used to track the animals in testing fields and to analyze the data.

### Open field test

Mice were placed into a white square open field chamber (45 × 45 × 40 cm, Panlab) under ambient light conditions, and activity was monitored for 5 mins in a single trial, as previously described^28^. The field was evenly divided into 16 virtual squares; the central four squares were defined as the center zone, while the remaining 12 squares along the walls were defined as the periphery. Mice were monitored by an overhead video camera and analyzed by SMART 3.0. Total distance traveled and the number of entries into the center zone were calculated.

### Novel object recognition

Novel object recognition was conducted as previously described^28, 95^ to assess the learning and memory with a 24-hour delay. Briefly, mice were habituated in a square open field chamber (45 × 45 × 40 cm) for two consecutive days, 10 min/day, under ambient light conditions. On the third day, two identical objects, termed “familiar” objects, were placed in the chamber, and mice were allowed to explore the objects freely for 10 minutes. After exploration, mice were returned to the home cage. After 24 hours, one of the “familiar” objects was replaced with a “novel” object. The mouse was again placed in the chamber and allowed to explore the objects for 10 mins. The total distance traveled, and the time spent exploring the objects were analyzed. The preference for the novel object was calculated as Time exploring novel object / (Time exploring novel object + Time exploring familiar object).

### Y maze

The spatial working memory of mice was assessed using a Y-maze as previously described^96^. A light-grey colored Y maze (30 × 6 × 15 cm, Panlab) with equal length arms was adopted in the study. Spontaneous alternation was calculated to evaluate the spatial working memory, since this behavior is driven by the innate curiosity of rodents to explore previously unvisited areas. The maze was divided into center zone and arms, which were labeled as zone A, zone B, and zone C, by SMART 3.0 software. The mouse was placed into the distal end of zone A and was allowed to explore the maze freely for 10 mins. A triplet entry into three different arms, e.g. from A to B then to C, was considered as one spontaneous alternation. Videos were taken during the test, and the percentage of alternation was calculated as (Number of Alternations / (Total number of arm entries-2)) × 100. Under a very rare condition, mice that failed to explore all three arms were excluded from downstream data analysis.

### Sample preparation and library construction for snRNA-seq, and sequencing

Chimeras from the control group (14 months old) and the P-tau 10 mpi group were used for single-nucleus RNA-sequencing. Three mice from each group were pooled into one sample, and two replicates were prepared for each condition (control and tau). Mice were euthanized and perfused with ice-cold DPBS, and the brains were quickly extracted and rinsed in the same buffer supplied with RiboGrip RNase inhibitor (Solis BioDyne, 0.8 U/μL). Brains were then dissociated and chopped into fine pieces before being transferred into Nuclei Extraction Buffer (Miltenyi Biotec, 130-128-024). Tissues were subsequently processed using gentleMACS dissociator (Miltenyi Biotec) following instructions. Extracted nuclei were filtered using 100 μm Smart Strainers (Miltenyi Biotec, 130-098-463) and then centrifuged at 300 × *g* for 5 mins at 4 °C. The pellet was resuspended using nuclei separation buffer and filtered again using 30 μm Smart Strainers (Miltenyi Biotec, 130-098-458). To obtain clean nuclei suspension with minimum cell debris, the suspension was treated with Anti-Nucleus Microbeads (Miltenyi Biotec, 130-132-997). After purification, nuclei were washed with PBS and proceed to library preparation using Chromium GEM-X Single Cell 3’ v4 Gene Expression kit (10XGenmoics) (Supplementary **Fig. S5A**). Quality controls of samples were performed by Genomics and Genome Editing Facility at Purdue University using High Sensitivity D5000 ScreenTape®. All samples passed quality control and were subsequently processed to downstream sequencing. snRNA-seq was performed by the Medical Genomics Core at Indiana University School of Medicine. 150bp paired-end data were generated on an Illumina NovaSeq X PLUS. The sequence reads were next processed using the 10X Genomics cellranger 9.0.1. The human plus mouse reference genomes (GRCh38_and_GRCm39-2024-A) were used for the alignment. The filtered feature-cell count matrices were used for further analysis.

### Single-nucleus RNA-sequencing data processing and quality control

Raw FASTQ files were processed using Cell Ranger (10x Genomics). Reads were aligned to a combined human–mouse reference, comprising of the human genome GRCh38 (GENCODE v32) and the mouse genome mm10 (GENCODE vM23). We labeled nuclei with >95% of reads mapping to the human or mouse reference genome as ‘human’ or ‘mouse’, respectively. This resulted in 25,230 human and 26,619 mouse nuclei prior to downstream filtering.

Quality control was performed using Seurat (v5.4.0). To set species-specific upper thresholds for gene detection and sequencing depth, we calculated the 99th percentile of nFeature_RNA and nCount_RNA distributions separately for human and mouse nuclei. Nuclei were retained if they had nFeature_RNA and nCount_RNA below the corresponding 99th percentile cutoffs and mitochondrial read content <15%. Following quality control filtering, 24,887 human and 26,288 mouse nuclei were retained. Doublets were subsequently identified and removed using the scDblFinder R package (v4.5.2), yielding final datasets of 22,784 human and 23,811 mouse nuclei (Supplementary **Fig. S5B, C**).

### Normalization, dimensionality reduction, clustering, and cell type annotation

Downstream analyses were performed using Seurat (v5.4.0). Gene expression counts were normalized and variance-stabilized using the SCTransform() function (vst.flavor = “v2”), which also identified the top 3,000 highly variable genes using default parameters. Principal component analysis (PCA) was performed on variable features, and a shared nearest-neighbor graph was constructed using FindNeighbors(). Cell clustering was carried out using FindClusters() at a resolution of 0.2. Human and mouse nuclei were annotated separately based on canonical marker genes for each cell type, as shown in dot plot visualizations.

### Within-species differential gene expression analysis

Differentially expressed genes (DEGs) were identified using Seurat’s FindMarkers() function with the Wilcoxon rank-sum test, with p-values adjusted for multiple testing using the Benjamini–Hochberg procedure and a minimum expression threshold of 10% of cells (min.pct = 0.1). Within-species differential expression was performed by comparing tau versus control nuclei within each annotated cell type. Genes were considered differentially expressed if they met both |avg_log2FC| ≥ 0.25 and adjusted p-value < 0.05 thresholds.

### Cross-species differential expression analysis

To enable cross-species comparisons, human gene symbols were mapped to mouse orthologs prior to differential expression analysis using the gprofiler2 R package (v4.5.2). Ortholog mapping was performed using the gorth()function, specifying ‘hsapiens’ as the source organism and ‘mmusculus’ as the target organism. Only one-to-one orthologs were retained for downstream cross-species analyses. Cross-species differential expression analysis was performed by comparing human and mouse nuclei within matched cell types following ortholog harmonization. Genes with an |avg_log2FC| ≥ 0.25 and adjusted p-value < 0.05 cut-off were considered differentially expressed. To assess condition-dependent species effects, a difference-in-differences (ΔΔlog2FC) metric was computed per gene and cell type. This metric provided the difference between the human–mouse log2 fold change in the tau condition and the corresponding human–mouse log2 fold change in the control condition. This ΔΔlog2FC metric was used to identify genes showing tau-associated divergence or convergence between species. ΔΔ log_2_ FCg,c = (log_2_ FCg,cHuman,Tau − log_2_ FCg,cMouse,Tau) – (log_2_ FCg,cHuman,Control − log_2_ FCg,cMouse,Control), g = gene; c = cell type. To visualize concordant and discordant tau-associated transcriptional responses between species, quadrant plots were generated by plotting tau-versus-control log2 fold changes in human neurons (human excitatory and inhibitory neurons) against corresponding log2 fold changes in mouse neurons (mouse excitatory and inhibitory neurons). Genes were categorized into four quadrants based on the direction of change in each species (Up–Up, Down–Down, Up–Down, Down–Up). Top genes per quadrant were prioritized based on combined effect size and highlighted for visualization.

### Pathway enrichment analysis

Functional enrichment analyses were performed on DEG sets using Reactome databases. Multiple hypothesis testing correction was applied using the Benjamini–Hochberg method.

### Statistics and reproducibility

Data are presented as mean ± SEM. All datasets were assessed for normality and were considered normally distributed. For comparisons between two groups, unpaired two-tailed Student’s t test was used. For comparisons among multiple groups, one-way ANOVA was performed. When variance was unequal across groups, variance-corrected tests were applied (e.g., Welch’s correction). A P value < 0.05 was considered statistically significant.

## Acknowledgments

This work was in part supported by grants from the NIH (R21NS140907 and U01AG088662 to R.X.), the Purdue Institute for Integrative Neuroscience research grant (iPSC/Organoid Technologies) to X.R, the Roberts AD Pre-Clinical Translational Science Grant to R.X., and the Showalter grant to R.X. Dr. Rochet’s lab is supported by NIH grants (R01AG098151 and R21NS135424). We thank Dr. Ying Liu (Florida International University) for providing the ND2, UTY-1 iPSC, and CAGG hESC lines; Dr. David Eliezer (Weill Cornell Medical College) for the pRK172-K18-P301L plasmid used to generate recombinant K18 P301L tau; and Dr. Marc Diamond (UT Southwestern Medical Center) for the HEK293T tau biosensor line. We also acknowledge Akhil Pinnapareddy, Oliver Thomas Johnson, and other members of the Xu Laboratory for their technical assistance and support.

## Authors’ contributions

R.X. and J.Y. designed the experiments and interpreted the data. J.Y. carried out most of the experiments with technical assistance from H.W., C.B., G.D., and Z.L., L.F. prepared the pathological tau seeds with quality control. C.B. prepared pH-rodo labeled K18 tau PFF with quality control. M.P., J. L. and J.Y. analyzed snRNA-seq data with help from L.X. and W.C. and instructions from B.P., and K.H. H.W., L.F., R. JC., Y. C., S.R., Q. C., and Y. Y. provided critical suggestions. R.X. directed the project and wrote the manuscript together with J.Y., with input from all co-authors.

---

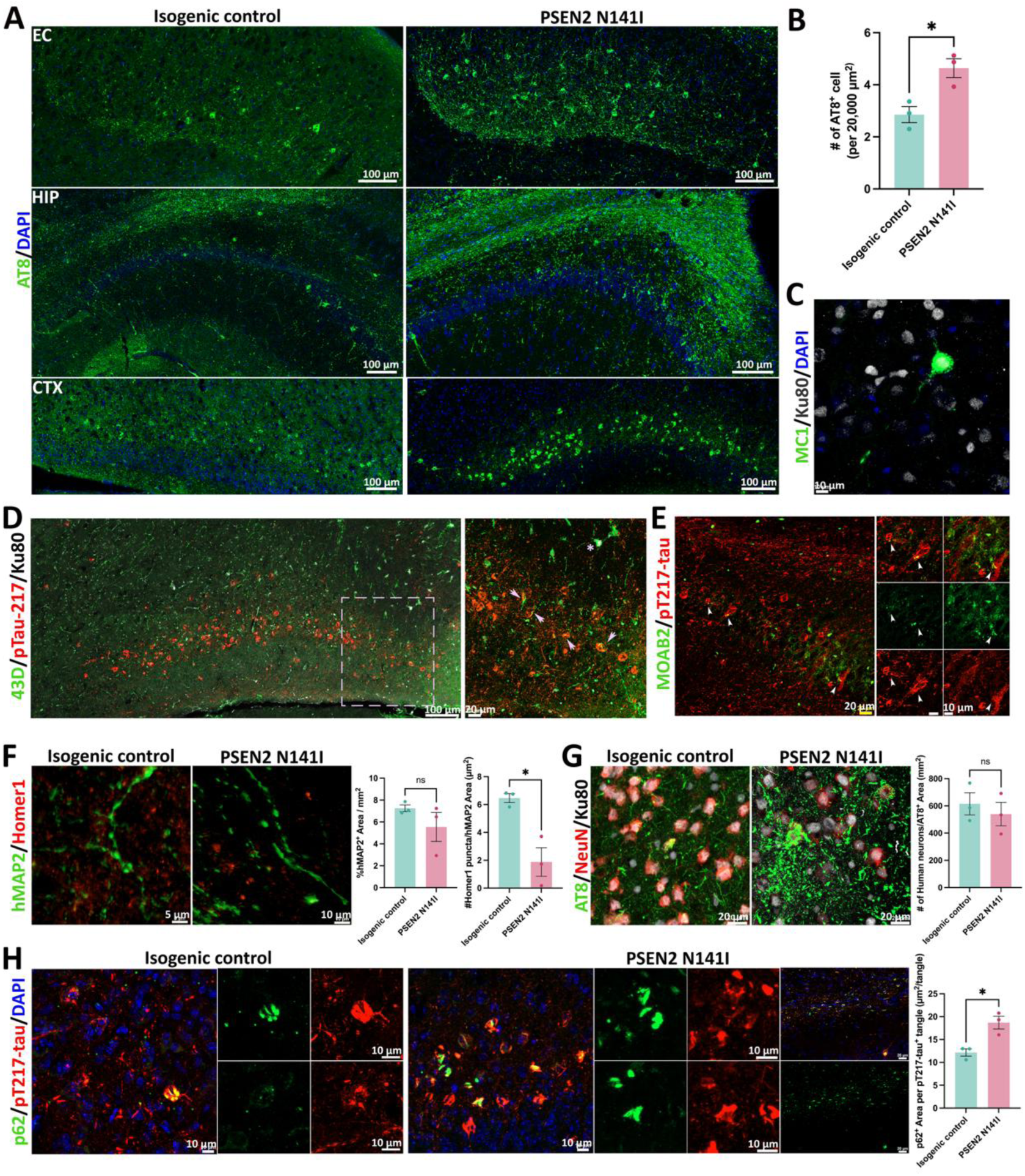

